# Structural characterization of the antimicrobial peptides myxinidin and WMR in bacterial membrane mimetic micelles and bicelles

**DOI:** 10.1101/2021.03.30.437760

**Authors:** Yevhen K. Cherniavskyi, Rosario Oliva, Marco Stellato, Pompea Del Vecchio, Stefania Galdiero, Annarita Falanga, Sonja A. Dames, D. Peter Tieleman

## Abstract

Antimicrobial peptides are a promising class of alternative antibiotics that interact selectively with negatively charged lipid bilayers. This paper presents the structural characterization of the antimicrobial peptides myxinidin and WMR associated with bacterial membrane mimetic micelles and bicelles by NMR, CD spectroscopy, and Molecular Dynamics simulations. Both peptides adopt a different conformation in the lipidic environment than in aqueous solution. The location of peptides in micelles and bicelles has been studied by paramagnetic relaxation enhancement experiments with paramagnetic tagged 5- and 16-doxyl stearic acid (5-/16-SASL). Multi-microsecond long molecular dynamics simulations of multiple copies of the peptides were used to gain an atomic level of detail on membrane-peptide and peptide-peptide interactions. Our results highlight an essential role of the negatively charged membrane mimetic in the structural stability of both myxinidin and WMR. The peptides localize predominantly in the membrane’s headgroup region and have a noticeable membrane thinning effect on the overall bilayer structure. Myxinidin and WMR show different tendency to self-aggregate, which is also influenced by the membrane composition (DOPE/DOPG versus DOPE/DOPG/CL) and can be related to the previously observed difference in the ability of the peptides to disrupt different types of model membranes.

## Introduction

Antimicrobial resistance represents a serious threat to global health, requiring urgent and concerted actions to fight a global crisis and the need to find alternative antimicrobial strategies^1–6^. Antimicrobial peptides (AMPs) are molecules widely distributed in nature which are rapidly gaining attention for their clinical potential and for their advantages compared to traditional antibiotics. AMPs are found in all forms of life, including bacteria, vertebrate, and invertebrate species^7–10^. Due to increasing resistance to currently used antibiotics, AMPs are promising candidates to build a new class of alternative broad-spectrum antibiotics^11^. Most AMPs are 12-50 amino acids long. Based on the physicochemical properties of AMPs and their target membranes, different mechanisms for their action have been described^12^. The cationic nature of AMPs arising due to a surplus of positively charged lysine or arginine residues compared to negatively charged glutamate and aspartate residues is critical for their selective action against bacterial membranes that contain negatively charged lipids like phosphatidylglycerol (PG) and cardiolipin^9,10,13–16^ and play a key role in the innate immune system. They are classified according to different criteria, but the most widely diffused classification is based on their secondary structure: α-helical, β-sheet, extended, and cyclic. Notwithstanding the differences in secondary structure, they all contain high amounts of arginine, tryptophan, histidine and glycine amino acids and carry net positive charge. The main mechanism of action is via direct interaction with the bacterial cell membrane, which is highly favored by the presence of i) positive charges for the initial interaction with the negatively charged bacterial membrane, ii) the presence of aromatic residues which are likely located at the interface between the membrane bilayer and the aqueous solution, iii) the ability to adopt an amphipathic structure in bacterial membranes. In particular, their net positive charge enhances electrostatic interactions between the cationic AMPs and anionic bacterial membranes stabilizing the binding, while the amphipathic structure leads to insertion of AMPs into the membranes, destabilization and disruption of the bacterial membrane. The hypothesized mechanisms of membrane disruption have been extensively reviewed^2,17,18^. Briefly, AMPs binding leads to a breakdown of membrane potential, an alteration in membrane permeability, causing bacterial cell death; in addition to their direct activity on the membrane bilayer, some AMPs have also an intracellular target.

Myxinidin is a marine peptide (NH_2_-GIHDILKYGKPS-CONH_2_ with a net charge of +2, Fig. 1A) isolated from the epidermal mucus of hagfish (*Myxine glutinosa L.*), which showed a significant antimicrobial activity against a wide range of bacteria and yeast and it demonstrated high levels of activity against *Pseudomonas aeruginosa* and *Escherichia coli* with low cytotoxicity against human cells^19,20^. A later modification of the myxinidin sequence led to the analogue WMR (NH_2_-WGIRRILKYGKRS-CONH_2_ with a net charge of +6, Fig. 1A), which has a higher antimicrobial activity compared to myxinidin against Gram positive and Gram negative bacteria^19,20^. In particular, WMR contains a tryptophan residue at the N-terminus, which usually is responsible for a strong membrane-disruptive activity and a higher number of positively charged amino-acids (arginines) compared to the native sequence. WMR has been exploited to obtain nanofibers which were shown to significantly inhibit biofilm formation and eradicate the already formed biofilms of *P. aeruginosa* (Gram-negative bacteria) and *Candida albicans* indicating that WMR-K is an interesting AMP to be further developed for its antimicrobial and antibiofilm activities^21^.

**Fig. 1:**
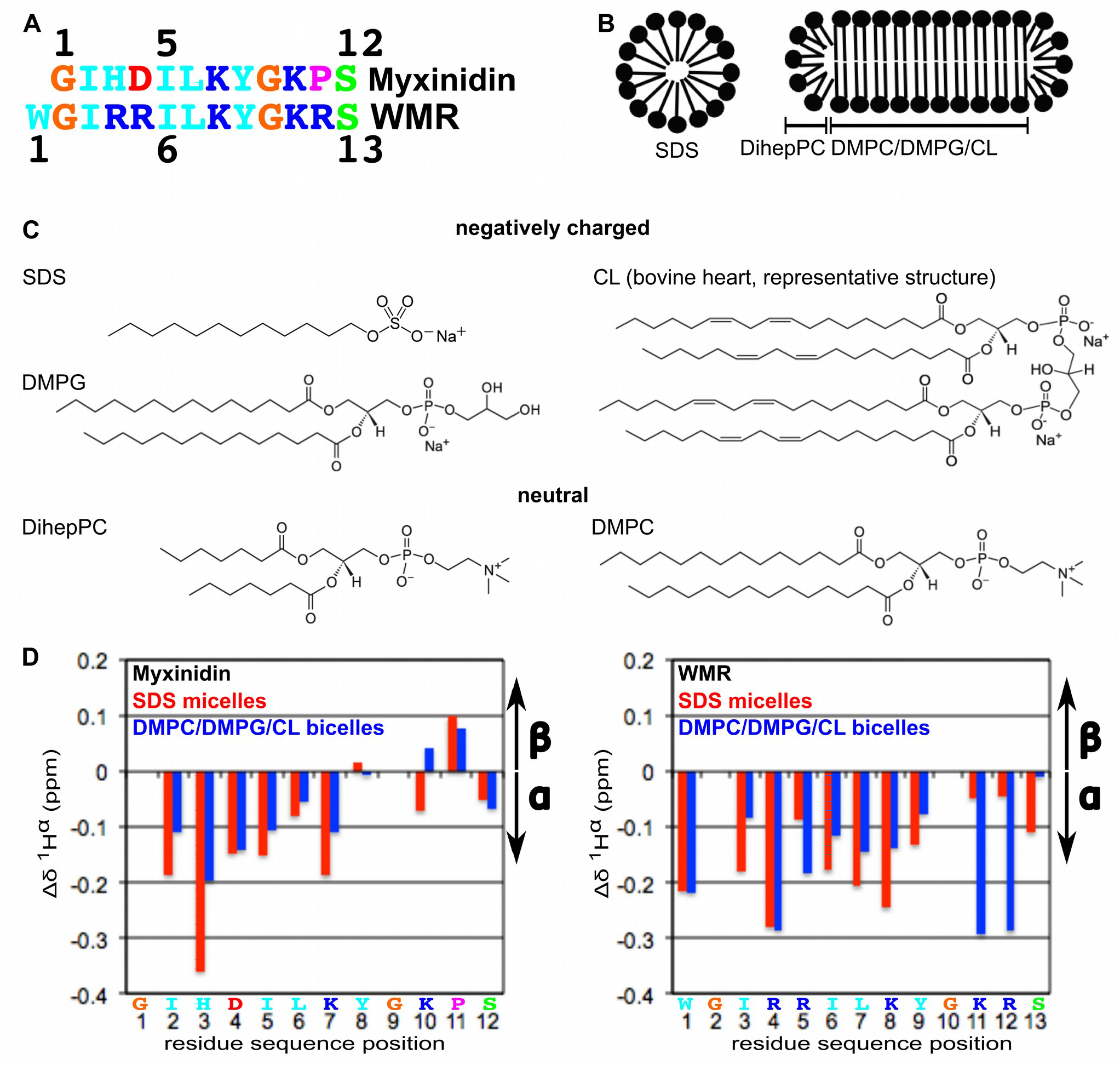
Primary and secondary structure of the antimicrobial peptides myxinidin and WMR and schematic representations of the used negatively charged membrane mimetics and their components. **(A)** Amino acid sequence of the 12-residue peptide myxinidin and a 13-residue long variant of it called WMR that has N-terminally an extra tryptophan and has arginines at positions 3, 4, and 11 of the original myxindin sequence. WMR has been shown to show higher antimicrobial activity^20^. Positively and negatively charged residues are colored in blue and red, respectively, aromatic and aliphatic residues in cyan, glycine in orange, serine in green and proline in magenta. **(B)** Schematic representation of a detergent micelle and a lipid bicelle. Negatively charged membrane mimetics can for example be obtained by forming micelles composed of sodium dodecyl sulfate (SDS) or by preparing the lipid bilayer of a bicelle from a mixture of the neutral phosphoslipid 1,2-dimyristoyl-*sn*-glycero-3-phosphocholine (DMPC) and the negatively charged phospholipids 1,2-dimyristoyl-*sn*-glycero-3-phospho-(1’-rac-glycerol) (DMPG) and cardiolipin (CL) (in this study 65:23:12 mole %) and the neutral short chain lipid 1,2-diheptanoyl-*sn*-glycero-3-phosphocholine (DihepPC) for the rim. **(C)** The chemical structures of the used membrane mimetic components of this study (from the website from Avanti Polar Lipids: https://avantilipids.com/). **(D)** The secondary structure content of myxinin (left) and WMR (right) in negatively charged SDS micelles (red bars) and DMPC/DMPG/CL bicelles (blue bars) as derived based on the measured ^1^Hα secondary shifts. Both peptides adopt a mostly helical structure upon membrane interactions, which is consistent with the CD data in Fig. 2 and SI Fig. S1.

Still the molecular mechanism underlying both myxinidin and WMR activities is not well understood. Although the disruption of the membrane bilayer has been demonstrated we cannot exclude the presence also of an intracellular target, thus more studies are needed. To better understand the molecular basis of the differences in the antimicrobial activity of myxinidin and WMR, we previously focused our interest in unraveling the mode of interaction with two different model bio-membranes, composed by DOPE/DOPG (80/20% mol) and DOPE/DOPG/CL (65/23/12% mol), mimicking respectively *E. coli* and *P. aeruginosa^22^* through a combined approach providing a comprehensive and detailed analysis of the peptide-membrane interactions, which clearly showed that the presence of CL lipid plays a key role in the WMR-membrane interaction.

To better understand the association of the natural AMP myxinidin and the more potent WMR, with different bacterial membrane mimetics, we analyzed their interaction with negatively charged membrane mimetic micelles and bicelles by NMR, CD spectroscopy, and molecular dynamics simulations. More information about the immersion properties in micelles and bicelles was derived from NMR studies using SDS micelle and DMPC/DMPG/cardiolipin bicelles containing paramagnetically tagged 5- and 16-doxyl stearic acid and in membranes from MD simulations. Molecular dynamics simulations of multiple copies of the peptides were used to gain an atomic level of detail on membrane-peptide and peptide-peptide interactions.

## Methods

### Peptide preparation and biophysical properties

Myxinidin (NH_2_-GIHDILKYGKPS-CONH_2_) and WMR (NH_2_-WGIRRILKYGKRS-CONH_2_) peptides were synthesized using the standard solid phase 9-fluorenylmethoxy carbonyl (Fmoc) method as previously reported^19^. The purified peptideCD spectra of myxinidin and WMR were performed by using a J-1500 spectropolarimeter (Jasco Analytical Instruments, Tokyo, Japan). Spectra were recorded in the 190 to 260 nm wavelength interval range, with 0.5 nm step resolution, 20 nm min^−1^ scan speed, 4 s response time, and 2 nm bandwidth, using a 0.1 cm path length quartz cuvette, at fixed temperature of 25°C. Cell cuvette thickness, peptide concentration and lipid concentration of vesicles were chosen in a way that the maximum high-tension voltage of the photomultiplier was not exceeding 600 V at the lowest wavelength (190 nm). Each experiment was reported as the average of 3 accumulated scans. The spectra were analyzed with JASCO software. For each sample, a background blank of either solvent or lipid vesicles without peptide was subtracted.s were obtained with a good yield (approximatively 60%) and identity was confirmed using a LTQ-XL Thermo Scientific linear ion trap mass spectrometer. The molar extinction coefficients that were determined spectroscopically by UV-Vis are ε (275nm) = 1647 ± 159 M^−1^ cm^−1^ for myxinidin and ε (280nm) = 4777 ± 281 M^−1^ cm^−1^ for WMR.

### Circular Dichroism measurements

CD spectra of myxinidin and WMR were performed by using a J-1500 spectropolarimeter (Jasco Analytical Instruments, Tokyo, Japan). Spectra were recorded in the 190 to 260 nm wavelength interval range, with 0.5 nm step resolution, 20 nm min^−1^ scan speed, 4 s response time, and 2 nm bandwidth, using a 0.1 cm path length quartz cuvette, at fixed temperature of 25°C. Cell cuvette thickness, peptide concentration and lipid concentration of vesicles were chosen in a way that the maximum high-tension voltage of the photomultiplier was not exceeding 600 V at the lowest wavelength (190 nm). Each experiment was reported as the average of 3 accumulated scans. The spectra were analyzed with JASCO software. For each sample, a background blank of either solvent or lipid vesicles without peptide was subtracted.

### Samples preparation for CD experiments

Liposomes with different composition were prepared: DOPE/DOPG (80/20 mole %) and DOPE/DOPG/CL (65/23/12 mole %). The lipids were weighted in a glass vial and dissolved in a chloroform/methanol mixture (2/1 v/v). A thin film was produced by evaporating the organic solvent with dry nitrogen gas. Lipid film samples were kept under vacuum for at least 4 h to remove the residual traces of the organic solvent. Dry lipids were then hydrated with 10 mM phosphate buffer, pH 7.4, vortexed obtaining multilamellar vesicles (MLVs), then sonicated obtaining the small unilamellar vesicles (SUVs). CD spectra of myxinidin were obtained by mixing a solution of the peptide with SUVs, at the lipid-to-peptide ratio of 20. The final peptide concentration was 50 μM. Due to WMR-induced vesicles aggregation, a different protocol for samples preparation was followed. Briefly, a solution of WMR peptide in 2,2,2-trifluoroethanol (TFE) was prepared. This solution was mixed with an equal volume of lipids (DOPE/DOPG or DOPE/DOPG/CL) dissolved in the organic solvent. Then, the solution was dried under gentle nitrogen steam and placed under vacuum to remove all the traces of solvent. The dry film was then hydrated with buffer solution (10 mM sodium phosphate, pH 7.4) to yield a final total lipid concentration of 1 mM and 50 μM of WMR peptide (L/P ratio of 20). Finally, the suspension was sonicated in order to obtain SUVs.

SDS micelles for the CD studies were formed by dissolving an appropriate amount of SDS in sodium phosphate buffer (10 mM, pH 7.4) to result in either a 20 or 100 mM stock solution. Deuterated d_25_-SDS micelles for the NMR studies were prepared by placing a defined amount of a 0.3 M stock solution of d_25_-SDS in chloroform/ethanol/water (65/35/8 v/v) in a glass vial and drying it under a stream of nitrogen gas. The dried SDS film was then dissolved in buffer and/or the protein sample to yield a final SDS concentration of 150 mM.

### Preparation of membrane mimetics for the NMR measurements

Sodium dodecyl sulfate (SDS) and cardiolipin from bovine heart were bought from Sigma Aldrich. 1,2-Dimyristoyl-*sn*-glycero-3-phatidylethanolamine (DMPE), 1,2-dimyristoyl-*sn*-glycero-3-phosphocholine (protonated = DMPC and deuterated = d_54_-DMPC), 1,2-dimyristoyl-*sn*-glycero-3-phospho-(1’-rac-glycerol) (DMPG), and 1,2-diheptanoyl-*sn*-glycero-3-phosphocholine (DiHepPC) were purchased from Avanti Polar Lipids.

For the samples with negatively charged bicelles (long chain lipids: DMPC/DMPG/cardiolipin 65/23/12 mole %, short chain lipid: DihepPC, q = 0.25, c_L_=11%) the appropriate amounts of stock solutions of the long chain lipids (DMPC or d_54_-DMPC, DMPG, cardiolipin) in chloroform were placed in a glass vial and dried under a stream of nitrogen gas. Bicelles were formed by stepwise addition of the appropriate amount of a DihepPC stock solution in buffer and vigorous vortexing after each step. Lastly, the protein solution was added and everything mixed by vortexing.

### Sample preparation for NMR experiments

For the samples used to record NMR data to assign and structurally characterize the peptides in the presence of negatively charged membrane mimetic micelles (150 mM d_25_-SDS) or bicelles (long chain lipids: d_54_-DMPC/DMPG/cardiolipin 65:23:12 mole %, short chain lipid: DihepPC, q = 0.25, c_L_=11%), the concentrations ranged of from 0.9 to 1.8 mM in PBS buffer (pH 7.4) supplemented with 0.02 % NaN3 and 10 % D_2_O (v/v).

### NMR spectroscopy

NMR spectra were acquired at 298 K on a Bruker Avance 500 MHz spectrometers equipped with a cryogenic probe. The data were processed with NMRPipe^23^ and analyzed using NMRView2^4^. Proton resonances were assigned based on two dimensional ^1^H-^1^H TOCSY, COSY, and NOESY experiments available in the Bruker standard pulse library (dipsi2esfbggph, cosycwgppsqf, and noesyfpgpphwg, respectively). The mixing time for the TOCSY experiments for the assignments as well as to obtain paramagnetic relaxation enhancement data of the peptides in membrane mimetics containing a spin label was 70 ms, only for WMR in bicelles it was 30 ms. The mixing time for the NOESY experiments was 100 ms and 200 ms. For the calculation of ^1^Hα secondary shifts, random coil values from the literature^25,26^ were subtracted from the measured chemical shifts.

### Structure calculations

All structure calculations were performed with XPLOR-NIH^27^ using molecular dynamics in torsion angle and Cartesian coordinate space. The amidated C-terminus of the peptides was taken into account by using the CTN option for the C-terminal residue for the generation of the psf file. Distance restraints were generated in NMRView and classified according to NOE-crosspeak intensities. Upper bounds were 2.8 Å, 3.5 Å, 4.5 Å, and 5.5 Å. The lower bound was always 1.8 Å. For all NOE-restraints r^−6^ sum averaging was used. For regions with α- or 3^10^- helical conformation, hydrogen bond restraints and for the structure calculations of the peptides in bicelles additionally backbone dihedral angle restraints for Φ and Ψ angles were derived based on the determined ^1^Hα chemical shifts and specific NOE-correlations^28^. Hydrogen bonds were defined by HN-O distance bounds of 1.8-2.3 Å, and N-O distance bounds of 2.6-3.1 Å. For the structure calculation of myxinidin in SDS micelles, initially two α-helix typical hydrogen bond restraints (i to i+4, i.e. 2 to 6 & 3 to 7) were used, however for the final run only NOE restraints were used. The spectra of WMR in the presence of SDS micelles showed more signal overlap than that of myxinidin. Interpretation of the spectra of both peptides in the presence of bicelles in the aliphatic region was challenging due to strong lipid signals. In these 3 cases the observed NOE correlations could not clearly discriminate between α- or 3^10^- helical conformation. Since the distortive lipid signals did generally hamper the detection of α-helix typical NOE cross peaks between the Hα of residue i and the Hβs of residues i+3, hydrogen bond restraints were used to support the helical structure indicated by the H^α^ chemical shifts (Fig. 1D), e.g. for Myxinidin in bicelles three hydrogen bond restraints for the region from I2 – Y8 were used. Since the NOE data did not allow to discriminate between α- and 3_10_-helical structure, we used ambiguous hydrogen bond restraints (i to i+3 or i+4, i.e., 2 to 6 or 5, 3 to 7 or 6 & 4 to 8 or 7). Backbone dihedral angle restraints for φ and ψ angles were restrained to values typical for helical regions (−65 ° ± 30 ° and −40 ° ± 30 °, respectively). The 20 lowest energy structures of in total 200 calculated structures were analyzed for the structural statistics and rendered with the software molmol^29^.

## Molecular dynamics simulations

### Myxinidin and WMR with SDS micelle

Molecular dynamics simulations of both myxinidin and WMR in an SDS micelle were performed with Gromacs 2016^30,31^ The peptide was initially placed at a random position near the preequilibrated micelle (75 SDS molecules) and solvated with ~32000 water molecules. First, Na^+^ ions were added to neutralize the system’s total charge, which was followed by the addition of Na^+^ and Cl^-^ ions to reach 0.1 M salt concentration. The CHARMM36m force field^32^ was used for the peptides and CHARMM36 for SDS. A 2 fs time step was used. All bonds were constrained with the LINCS algorithm^33^. Water bond lengths and angles were kept constant with the SETTLE algorithm^34^. Initial velocities were taken from the Maxwell distribution for 303.15 K. A constant temperature of 303.15 K was maintained with a V-rescale thermostat^35^ with 0.1 ps coupling constant. SDS micelle, peptide, and water with ions were coupled to separate thermostats with the same parameters. The constant pressure of 1 bar was maintained with an isotropic Parrinello-Rahman barostat^36^ with 5.0 ps coupling constant and a compressibility of 4.5×10^−5^ bar^−1^. The particle mesh Ewald algorithm^37,38^was used for long-range contributions to electrostatic interactions. Lennard-Jones interactions were cutoff at 1.2 nm, with a force-switch modifier from 1.0 to 1.2 nm.

Each system was equilibrated for 10 ns, followed by 500 ns of production run. Both peptides bound to the micelle within the first 5 ns of the production run, but the first 100 ns of the run were not used for analysis purposes to allow the peptide to fully equilibrate in a micelle-bound state. The distance between the micelle center of mass (COM) and separate peptide residues COM was computed. To analyze peptide stability, the secondary structure of each peptide was computed as a function of time with the gmx do_dssp analysis program, a part of the Gromacs package. The micelle surface for the images of the micelle-bound peptides was defined as an isosurface of averaged SDS density.

### Myxinidin and WMR with DOPE/DOPG, DOPE/DOPG/CL bilayers

Molecular dynamics simulations were performed with Gromacs 2016.3 ^30,31^. Six different systems were simulated: 18 myxinidin peptides with a DOPE/DOPG bilayer, 18 myxinidin peptides with a DOPE/DOPG/CL bilayer, 18 WMR peptides with a DOPE/DOPG bilayer, 18 WMR peptides with a DOPE/DOPG/CL bilayer, as well as DOPE/DOPG and DOPE/DOPG/CL bilayers without peptides as control. We also initially performed our simulation in the presence of a DMPC/DMPG bilayer, but this bilayer composition turned out to be unstable at 303 K with the CHARMM36 force field. The DMPC/DMPG membrane exhibited a spontaneous transition from liquid to interdigitated gel phase after a few microseconds of simulation, even without any peptides present. As a result, we do not present a detailed analysis of this simulation setup.

The DOPE/DOPG bilayer was composed of 144 DOPE lipid molecules and 36 DOPG lipid molecules (80/20 ratio). The DOPE/DOPG/CL bilayer was composed of 116 DOPE, 42 DOPG, and 22 cardiolipin molecules (18:2,18:2/18:2,18:2 lipid tails) with 65/23/12 ratio or also mole %. Initial conformations were generated by placing 18 copies of a peptide around the preequilibrated membrane at random positions. Next, each system was solvated with ~27000 water molecules, and Na^+^ and Cl^-^ ions were added to reach 0.1 M salt concentration. The CHARMM36m force field was used for the peptides and CHARMM36 for lipids. The same run parameters were used as in the simulations with SDS micelles unless otherwise noted.

To mimic the physiological situation, in which the peptides first can access only one side of the membrane, an additional flat-bottom potential was applied in the direction perpendicular to the membrane plane, between bilayer COM and the peptide backbone atoms to prevent peptides from accessing both sides of the membrane through periodic boundary conditions. This potential was different from zero if the distance between peptide and bilayer COM is greater than 7 nm. A force constant of 500 kJ/mol was used. As a result, peptides in the membrane-bound state were unaffected by the flat-bottom potential. Only detached peptides in the bulk solution were affected. The size of the systems in the z-direction (perpendicular to the membrane plane) fluctuated around 17-18 nm during the simulations. Each system was simulated for 5 microseconds. Analyses were performed on the last 2.5 microseconds of a trajectory. The results of bilayer-peptide simulations were compared to the corresponding bilayer only simulations. The tendency of peptides to form aggregates was estimated by calculating the probability that a randomly selected peptide will belong to an aggregate of size 1 (no aggregation) to 18 (all the peptide copies form a single aggregate). Two peptides were considered to belong to the same aggregate if they have contacts within 0.3 nm.

## Results

### CD data and ^1^Hα secondary shifts indicate that myxinidin and WMR adopt a more ordered, rather helical structure upon interaction with negatively charged membrane mimetics

As SUVs and other vesicles are too large for NMR structure determination, we turned to micelles composed of negatively charged SDS or bicelles composed of DMPC, DMPG, and cardiolipin (65/23/12 mole %) as long-chain lipids and DihepPC as short-chain lipid (q = 0.25, cL 11% w/v) for the NMR structural charactrerization of myxinidin and WMR (Fig. 1B-C). Whereas the micelles could be prepared using fully deuterated SDS (d_25_), the bicelles were prepared using only deuterated DMPC (d_54_) but fully protonated DMPG and cardiolipin as well as DihepPC. Because of this and the smaller size of a micelle compared to a bicelle and thus a shorter rotational correlation time, the homonuclear ^1^H-^1^H TOCSY and NOESY data recorded for the assignment and to obtain distance restraints for structure determination showed less distortive signals in the presence of membrane mimetic SDS micelles compared to the DMPC/DMPG/CL/DihePC bicelles (SI Fig. S2-S5). Comparing the data for myxinidin (SI Fig. S2 in the presence of SDS bicelles and S3 in the presence of DMPC/DMPG/CL bicelles) and WMR (SI Fig. S4 in the presence of SDS bicelles and S5 in the presence of DMPC/DMPG/CL bicelles), the myxinidin spectra showed overall a much better signal dispersion and less signal overlap. This can be explained by the greater variability of the amino acid composition of the sequence of myxinidin compared to WMR (Fig. 1A). In the case of myxinidin, almost all backbone and side-chain protons could be assigned (see labels in SI Fig. S2 and S3 and SI table S1), including the protons of the C-terminal amide group (S12 H1 and H2). In the case of WMR, most ^1^H signals could be assigned in SDS micelles (SI Fig. S4, SI table S1). However, some side-chain protons of the arginine, isoleucine, and leucine residues could not be assigned in the presence of DMPC/DMPG/CL/DihepPC bicelles due to signal overlap and strong bicelle signals in the aliphatic region (SI Fig. S5, SI table S1).

Fig. 1D shows the ^1^H^α^ secondary chemical shifts of myxinidin and WMR in both membrane mimetics. Since they are negative for most residues, these data indicate that both peptides adopt a mostly α-helical structure in the presence of negatively charged SDS micelles and DMPC/DMPC/CL bicelles. This is further supported by the CD data of both peptides with the respective membrane mimetics shown in SI Fig. S1. Myxinidin is α-helical from I2 to K7 based on the ^1^Hα secondary shifts. Y8 preceding a glycine shows no specific preference. K10 preceding the proline shows a more typical α-helix shift in micelles and a more β-sheet like one in bicelles. Glycines increase the local flexibility and allow due to their small size for kinks or loops in the backbone, and prolines have been shown to locally restrict the backbone conformation^39^. Thus, the C-terminus may still form a turn-like structure. WMR shows an α-helical secondary structure for W1-S12 and thus the whole peptide. However, whereas the helical character for I3 and I6-Y9 is higher in SDS micelles, it is higher for R4 and even more for K11 and R12 in DMPC/DMPG/CL bicelles. Note that no ^1^H^α^ secondary shift is given for glycines because it has two α-protons (G1 and G9 in myxinidin and G2 and G10 in WMR). Based on the ^1^H^α^ secondary shifts, myxinidin and WMR in negatively charged micelles and bicelles adopt a mostly helical structure (Fig. 1D).

The estimate of the secondary structure content from the NMR data is in line with the CD data of myxinidin and WMR in the absence and presence of small unilamellar vesicles (SUVs) composed of DOPE/DOPG and DOPE/DOPG/CL, mimicking the plasma membrane of *E. coli*. and *P. aeruginosa* (Fig. 2) and negatively charged SDS micelles (SI Fig. S1). In the absence of membrane mimetics, both the myxinidin and WMR spectra (black) show a large negative band at about 200 nm, indicating that they are mainly not structured in buffer solution. In the presence of SDS micelles and DOPE/DOPG and DOPE/DOPG/CL vesicles, dramatic changes in the CD spectra were observed. In particular, for myxinidin in the presence of DOPE/DOPG vesicles (blue) two separated negative bands at around 205 nm and 220 nm were detected indicating that the peptide is adopting a helical structure. These general features were also observed in the presence of DOPE/DOPG/CL vesicles (cyan) where the negative bands are shifted towards longer wavelengths (208 nm and 222 nm) suggesting a more ordered structure in the presence of CL-containing vesicles. For the WMR peptide two distinct negative bands around 207 nm and 222 nm were detected in the presence of both DOPE/DOPG (blue) and DOPE/DOPG/CL (cyan) vesicles, showing that it is also able to adopt an ordered helical structure. The spectra changes in the presence of SDS micelles are both peptides very similar to those observed in the presence of SUVs.

**Fig. 2:**
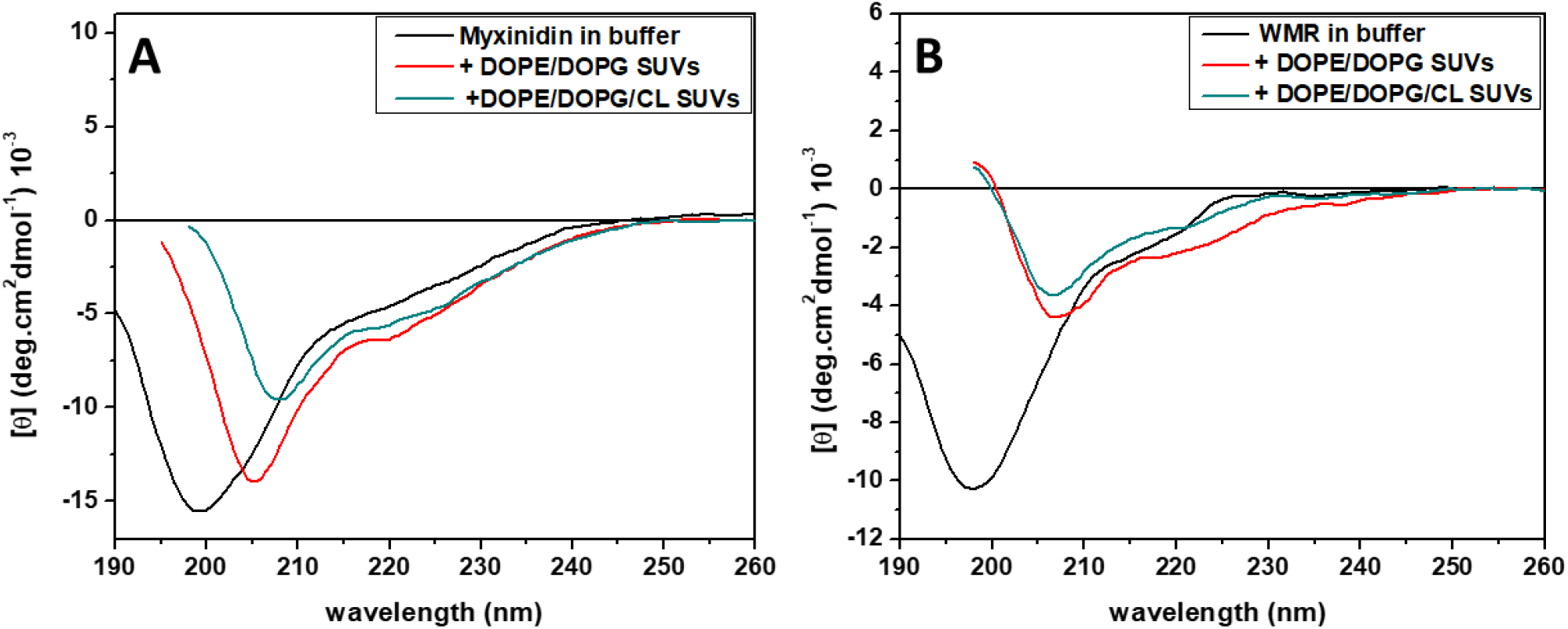
Far-UV CD spectra for myxinidin (A) and WMR (B) peptides in the buffer solution (black lines), in the presence of DOPE/DOPG vesicles OR SUVs (see comment in methods section) (blue lines) and in the presence of DOPE/DOPG/CL vesicles (cyan lines) at a lipid-to-peptide ratio of 20. All the spectra were recorded in 10 mM phosphate buffer, pH 7.4, at the temperature of 25 °C.

### Myxinidin adopts an amphipatic helical structure in the presence of negatively charged membrane mimetics

The three-dimensional structures of myxinidin and WMR in negatively charged membrane mimetic SDS micelles and DMPC/DMPG/CL bicelles (Fig. 3) were calculated based on distance restraints derived from the 2D ^1^H-^1^H NOESY and only if needed additional hydrogen bonds and/or backbone dihedral angle restraints. The structural statistics are given in table 1.

**Fig. 3:**
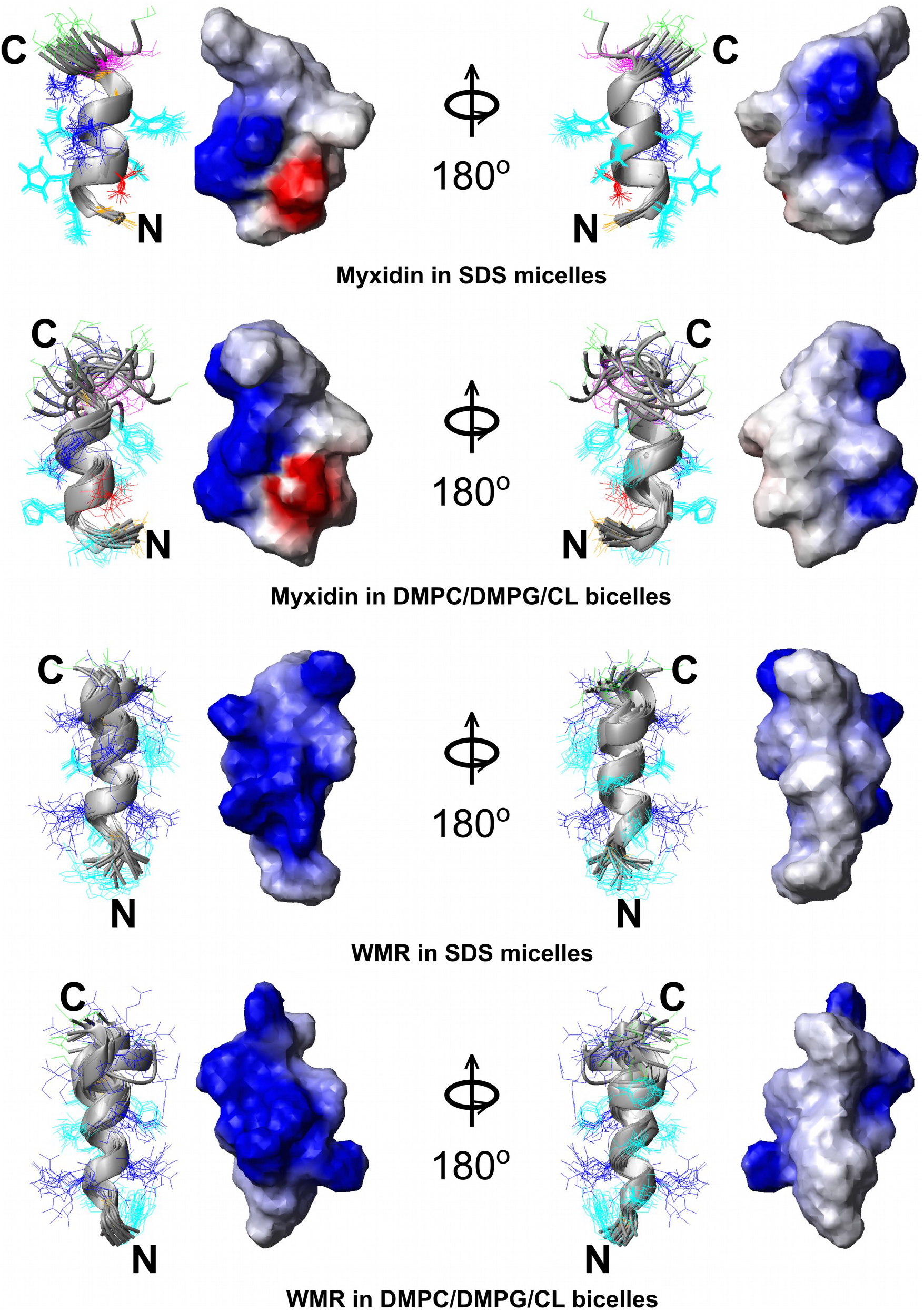
The three-dimensional structures of myxinidin and WMR in negatively charged membrane mimetics micelles and bicelles that have been calculated based on homonuclear ^1^H-NMR data. The top two panels show the structures of myxinidin in SDS micelles and DMPC/DMPG/CL bicelles, respectively and the bottom two those of WMR in the same membrane mimetics. In each plot half, a ribbon representation of a superposition of the 20 lowest energy structures is shown. The ribbon of the α- and 3^10^- helical regions is colored grey. The color coding of the side chain that are shown as line representations is the same as in Fig. 1A (cyan: aliphatic and aromatic, red: negatively charged, blue: positively charged, orange: glycine, magenta: proline, green: serine). The right half of each plot half shows a surface charge representation of the lowest energy structure (red: negatively charged, blue: positively charged). The structural statistics are given in table 1. In each horizontal plot, the right half represents the view after a 180° rotation around the vertical axis.

**Table 1.** Statistics for the 20 final structures of myxinidin or WMR bound to negatively charged membrane mimetics*

| Peptide, membrane mimetic | myxinidin, SDS micelles | myxinidin, DMPC/DMPG/ cardiolipin bicelles | WMR, SDS micelles | WMR, DMPC/DMPG/ cardiolipin bicelles |
| --- | --- | --- | --- | --- |
| Distance restraints | All (assigned + ambiguous) | All (assigned + ambiguous) | All (assigned + ambiguous) | All (assigned + ambiguous) |
| Total | 372 (331 + 41) | 207 (184 + 23) | 248 (215 + 33) | 171 (141 + 30) |
| NOESY 100 ms | 0 (0 + 0) | 0 (0 + 0) | 0 (0 + 0) | 17 (15 + 2) |
| NOESY 200 ms | 372 (331 + 41) | 204 (181 + 23) | 241 (208 + 33) | 149 (119 + 28) |
| Hydrogen bond restraints | 0 | 3 | 7 | 7 |
| $\phi + \psi$ angle restraints | 0 | 0 | 18 | 18 |
| Rmsd's from experimental restraints |  |  |  |  |
| Distance (Å) | 0.0289±0.0011 | 0.0298±0.0046 | 0.0185±0.0035 | 0.0190±0.0049 |
| Dihedral angle (°) | - | - | 0.106±0.089 | 0.100±0.090 |
| Rmsd's from idealized geometry |  |  |  |  |
| Bonds | 0.0116 ±0.0001 | 0.0114±0.0001 | 0.0040±0.0003 | 0.0029±0.0004 |
| Angles | 1.70±0.009 | 1.666±0.013 | 0.402±0.032 | 0.308±0.039 |
| Improper | 5.009±0.003 | 5.010±0.010 | 0.217±0.022 | 0.220±0.034 |
| Lennard Jones Energy (kcal mol<sup>-1</sup>) | -29.2±2.6 | -14.3±6.6 | -23.9±27.5 | -20.8±7.1 |
| Average rmsd to mean structure ( bb/heavy, Å) |  |  |  |  |
| Residues 1-12 | 0.56 / 0.87 | 1.91 / 3.08 | - / - | - / - |
| Residues 1-13 | - / - | - / - | 1.18 / 2.06 | 1.31 / 2.70 |
| Residues 2-9 | 0.22 / 0.65 | 0.66 / 1.25 | 0.63 / 1.58 | 0.61 / 2.14 |
| *None of the structures had distance restraints violations > 0.5 Å or dihedral angle violations > 5°. rms(d) = root mean square (deviation), bb = backbone |  |  |  |  |

For myxinidin in SDS micelles, about 370 NOEs could be assigned because of the good signal dispersion and small spectral distortions from the deuterated SDS and other buffer components. Consistent with the high number of distance restraints, the structure is overall very well defined and shows rmsd values of 0.22 Å for the backbone of residues 2-9 and 0.56 Å for the full sequence (residues 1-12). The C-terminal end encompassing K10-P11-S12 is overall less well defined compared to the α-helical stretch from residue I3-G9 that may extend to I2, which shows a turn-like secondary structure. The glycine at position 9 and the proline at position 11 presumably enable the C-terminus to bend back to the helical region. Consistent with the high number of distance restraints and the low backbone rmsd the side-chain conformations of residues 2-9 are also very well defined, which is reflected in a rmsd for all heavy atoms of 0.65 Å. The surface of myxinidin is amphipathic (Fig. 3 top panel) with a rather large hydrophobic patch at the mostly helical fold due to the aliphatic and aromatic residues at positions 2, 3, 5, 6, 8, and 11, a positive patch formed by K7 and K10 and a smaller negative one due to presence of D4. Based on the analysis of interatomic distances (SI results), the helical structure of myxinidin in the presence of micelles appears to be stabilized by a salt bridge interaction between D4 and the N-terminus and if at all a cation-π interactions between H3 and K7. Please note, that these interactions were not restrained by the NMR data. Whereas the hydrophobic side chains of the aromatic and aliphatic side chain may immerse the SDS micelle to make contacts with the hydrophobic acyl chains, the positively charged side chains of K7 and K10 may interact with the negatively charged sulfate groups of SDS.

The NMR structure of myxinidin in DMPC/DMPG/CL-DihepPC bicelles (Fig. 3, second panel) is overall rather similar to that in SDS micelles (Fig. 3, first panel). Due to the strong remaining signals from the undeuterated lipid components, especially in the aliphatic region, only 204 NOE restraints (Table 1) could be extracted from the 2D ^1^H-^1^H NOESY data (SI Fig. S3). Thus, the structural quality is lower, and the structure is overall less well defined, which is reflected in higher backbone and side chain rmsd values for the mostly helical region around residues 2-9 (0.66 and 1.25 Å, respectively, Table 1) and even higher ones if the C-terminal region around P10 is included (1.91 and 3.08 Å, respectively, table 1). The lower quality of the structure of myxinidin in bicelles compared to micelles is also reflected in a less negative average Lennard-Jones energy value and a higher standard deviation for the ensemble of the 20 lowest energy structures (Table 1). Due to the strong remaining signals from the undeuterated lipid components, especially in the aliphatic region, only 204 NOE restraints (Table 1) could be extracted from the 2D ^1^H-^1^H NOESY data (SI Fig. S3). Since the distortive lipid signals did generally hamper the detection of α-helix typical NOE cross peaks between the Hα of residue i and the Hβs of residues i+3, three hydrogen bond restraints for the region from I2 – Y8 were used to support the helical structure indicated by the H^α^ chemical shifts (Fig. 1D). Since the NOE data did not allow to discriminate between α- and 3_10_-helical structure, we used ambiguous hydrogen bond restraints (i to i+3 or i+4, i.e., 2 to 6 or 5, 3 to 7 or 6 & 4 to 8 or 7). In the 20 lowest energy structures of myxinidin in the presence of negatively charged bicelles, residues 3-8, in some structures even residues 2-8, adopt an α-helical conformation. The structure is similarly amphipathic as in micelles (Fig. 3, second panel). Consistent with the similarity to the structure in the presence of micelles that in the presence of bicelles may also be stabilized by ionic interaction between D4 and the N-terminus as well as by cation-π interactions between H3 and K7 (SI results). Whereas the hydrophobic side chains of the aromatic and aliphatic residues may be immersed in the membrane to make contacts with the hydrophobic lipid acyl chains, the charged residues may interact with the polar headgroups. The two lysines may thereby contribute to the increased affinity for negatively charged lipid bilayers.

### Based on NMR PRE and chemical shift mapping data the helical structure of myxinidin does not deeply penetrate negatively charged micelles or bicelles

In order to better understand how myxinidin associates with negatively charged membranes, we looked at the paramagnetic relaxation enhancement (PRE) and chemical shift changes of myxinidin in the presence of SDS micelles doped with stearic acid molecules containing a paramagnetic nitroxide group at position 5 or 16 of the acyl chain to which we refer to as 5- and 16-SASL. Based on former studies by ourselves and the literature, both spin labels reside rather close to lipid head group^40^. In the case of 16-SASL because the acyl chain bends, presumably because it is energetically more favorable to place the polar nitroxide group closer to the head groups than deep in the hydrophobic interior of the micelle^41^. Since we had only unlabeled peptides at hand, we recorded 2D ^1^H-^1^H TOCSY spectra in the presence of SDS micelles or DMPC/DMPG/CL/DihepPC bicelles without and with 5- and/or 16-SASL (SI Fig. S6-S9) and looked at the HN-H^α^ correlation of each residue. Since the spectra of myxinidin show generally a good signal dispersion, the reduction in signal intensity due to the PRE effect and the change of the chemical shift due to the change in the chemical environment between pure membrane mimetics and such doped with 5- and 16-SASL could be determined (Fig. 4). Generally, residues close to the lipid or detergent head groups should experience strong PRE effects, whereas residues deeper in the membrane mimetic or at the surface should experience weaker ones. Since the doxyl group does not induce pseudo contact shifts, the observed chemical shift changes reflect the change in the chemical environment between pure micelles and bicelles and such doped with 5- or 16-SASL. Myxinidin in micelles with 5-SASL shows the strongest PREs for residues I2, and K7 and a bit weaker ones for L6, Y8, G9, and K10 (Fig. 4B grey bars). This is similarly reflected in the spectral changes visible for the side chains (SI Fig. S6A, top). In contrast to those of the backbone, the side-chain resonances of H3 and I5 also show strong changes. The spectral changes with 16-SASL are overall stronger (Fig. 4A). This has similarly been observed for other proteins/membrane mimetic systems^42^. The correlations of the H^N^ of K7 to its H^α^ and side-chain proteins are broadened beyond detection with only 1.5 mM 16-SASL and those of I2 and K10 at 2.7 mM 16-SASL. Those of Y8 and L6 are very weak and those of H3 and I5 are significantly weakened at 2.7 mM 16-SASL. This suggests that all these residues are relatively close to the head group region. S12, G9, and D4 show rather small spectral changes, which suggest that they are more solvent-exposed. The data of myxinidin in bicelles with 5-SASL (SI Fig. S7) shows the strongest spectral changes also for the H^N^ to side-chain proton correlations of K7 and/or I5, which are overlapped in these data, as well as for L6 and I3. H3 and Y8 show weaker changes. Again S12, G9, and also D4 show only very weak to weak changes. Overall the data suggest that the helical structure of myxinidin immerses the bilayer mostly in the headgroup region and the nearby hydrophobic interior but does not penetrate it deeper.

**Fig. 4:**
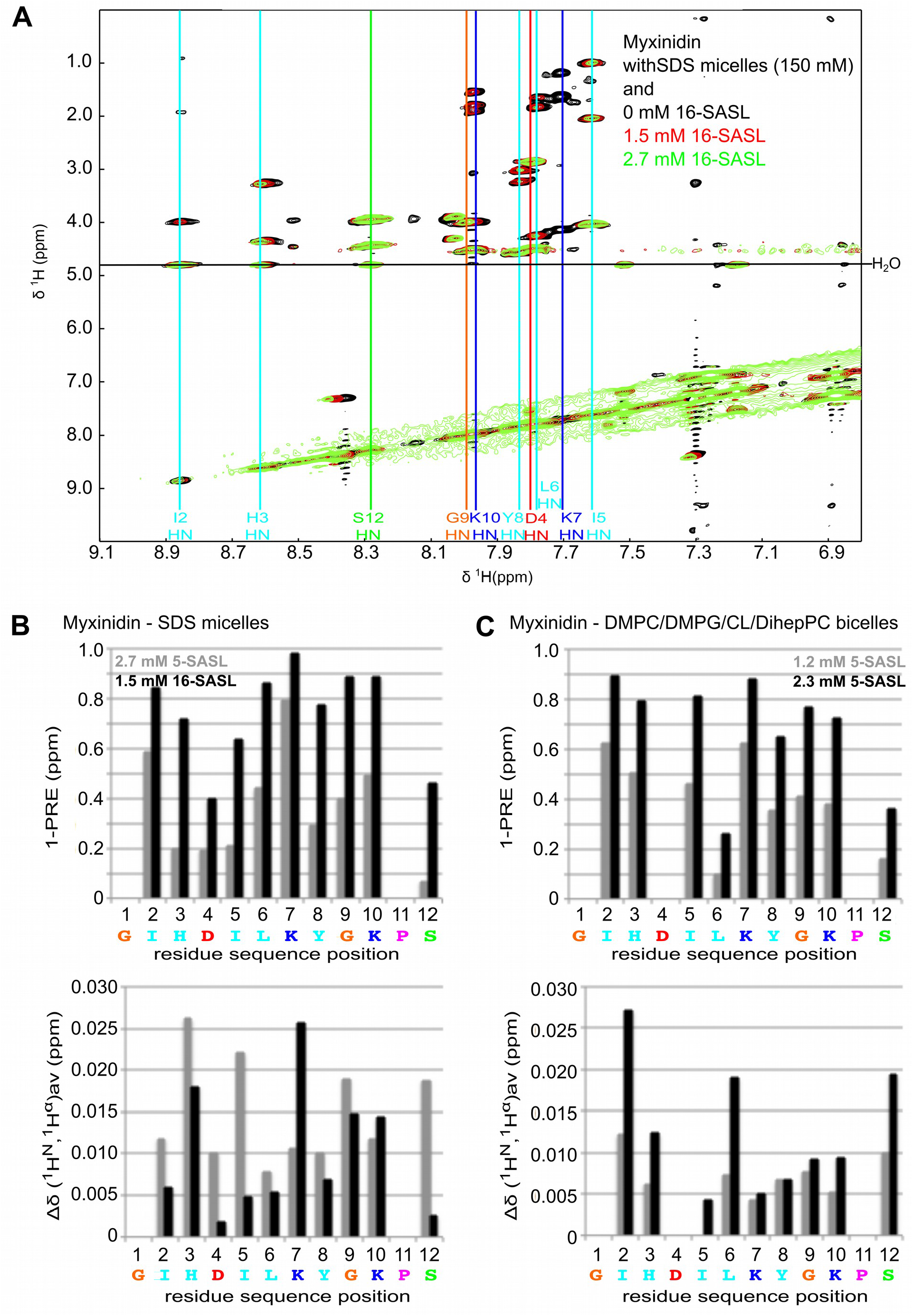
Analysis of the spectral changes of myxinidin in the presence of SDS micelles or DMPC/DMPG/CL bicelles doped with paramagnetic doxyl-labeled stearic acid molecules. A) shows the ^1^H-^1^H-TOCSY of myxinidin in the presence of SDS micelles with increasing concentrations of 16-SASL, which results in a reduction of the peak intensity due to paramagentic relaxation enhancement and/or a change of the chemical shift position due to a change in the chemical environment. The assigned amide and aromatic protons are labeled with the one-letter amino acid code, the residue sequence position and the atom name. The 2D ^1^H-^1^H-TOCSY spectrum in the presence of SDS micelles and 5-SASL is shown in SI Fig. S6 and that in the presence of DMPC/DMPG/CL bilayers and 5-SASL in S7. The data for WMR in the presence of 5- or 16-SASL in SDS micelles and 5-SASL in DMPC/DMPG/CL bicelles (SI Figs. S8-S9) could not be analyzed because the changes have been too strong and/or because of too much signal overlap. B, C) Shown are diagrams of the PRE effects and chemical shift changes of the H^N^-H^α^ cross peaks of membrane mimetic associated myxinidin in the presence of the indicated amount of 5-SASL as a function of the residue sequence position. To better compare the PRE effects to the chemical shift changes, 1-PRE (= 1- I(x mM SASL)/I (0 mM SASL)) was plotted. Accordingly, the larger the PRE effect, the higher the 1-PRE value. The sequence is given at the bottom.

### WMR adopts a largely α-helical structure in the presence of negatively charged membrane mimetics that is positively charged on one side and hydrophobic on the other

As for myxinidin, the structures of WMR in the presence of negatively charged SDS micelles and DMPC/DMPG/CL/DihepPC bicelles (Fig. 3 bottom two panels) are very similar. Both membrane mimetics induce a predominantly helical structure. The structural statistics are given in the third and fourth columns of Table 1. Again, the calculated structures for the micelle-associated state are better defined than for the bicelle-associated one, because the fully deuterated d_25_-SDS results in significantly fewer distortive signals than the only partially deuterated d54-DMPC/DMPG/CL/DihepPC bicelles. Thus, 241 NOE distance restraints (Table 1) could be extracted for WMR in the presence of micelles but only 166 in the presence of bicelles. However, due to the lower variation in the sequence composition of WMR (Fig. 1A) and the resulting lower signal dispersion of WMR (SI Fig. S4-S5) compared to myxinidin (Fig. 1A, SI Fig. S2-S3), the number of extracted distance restraints is overall lower than for myxinidin. Because of this, additional hydrogen bond and dihedral angle restraints were used for the region, which is based on the ^1^Hα secondary shift (Fig. 1D) helical (residues 3-13). The rmsd values for the ensemble of the 20 lowest energy structures for residues 2-9 are 0.63 / 1.58 Å (backbone/heavy atoms) for the micelle- and 0.61 / 2.14 Å for the bicelle-associated structures and for residues 1-13 1.18 / 2.06 Å (backbone/heavy atoms) and 1.31 / 2.70 Å, respectively. Based on the analysis of interatomic distances (SI results), the helical structure of WMR in the presence of micelles and bicelles could be stabilized by cation-π interactions between W1 and R5 as well as between Y9 and R12 as well as R5. Consistent with the presence of 5 positively charged residues in the 13-residue long sequence of WMR (Fig. 1A), about half of the surface of the helical structures in the presence of micelles and bicelles is positively charged, whereas the remaining half is hydrophobic (Fig. 3, bottom two panels). The large positively charged region can drive the initial interaction with the surface of negatively charged membranes, whereas the hydrophobic region may interact with the lipid acyl chains following a subsequent deeper immersion in the bilayer. As for myxinidin, we also recorded 2D ^1^H-^1^H TOCSY data of WMR in micelles and bicelles in the presence of 5- or 16-SASL (SI Fig. S8 and S9). However, due to significant signal overlap and in the case of micelles with 16-SASL due to very strong PRE effects and in the case of bicelles with 5-SASL also due to strong ridges, the data could not be interpreted quantitatively. As for myxinidin, the reduction of the signal intensity, and thus the PRE effects are stronger with 1.5 mM 16- than with 2.7 mM 5-SASL in SDS micelles (SI Fig. S8). Based on the data with 16-SASL (SI Fig. S8, bottom part), most residues show a strong PRE effect and thus appear to reside around the SDS head groups. Only the Hβ protons of the C-terminal serine show still a strong signal in the presence of 16-SASL in SDS micelles and may thus be more solvent exposed.

### MD simulation of myxinidin and WMR in the presence of a SDS micelle

To further investigate the interactions of myxinidin and WMR with SDS micelles on a molecular level, MD simulations of micelle-peptide complexes were performed. The peptides’ initial structure was taken as top 1 structure from the ensemble calculated based on the NMR data of myxinidin or WMR in SDS micelles. A single copy of myxinidin or WMR was placed in the water near the SDS micelle at the beginning of simulation. Myxinidin and WMR bind to the SDS micelle during the first few nanoseconds of the simulation and stay in a micelle-associated state for the whole duration of the 500 ns simulation (Fig. 5A). We performed a secondary structure analysis of myxinidin and WMR as a function of time to monitor the peptide structure in the SDS micelle-associated state (SI Fig. S10). Both peptides keep their mostly α-helical structure during the whole duration of the simulation, with WMR exhibiting slightly higher variability in secondary structure. This result is in line with the experimental findings presented in the current paper that negatively charged SDS micelle stabilize an α-helical structure of myxinidin and WMR peptides.

**Fig. 5:**
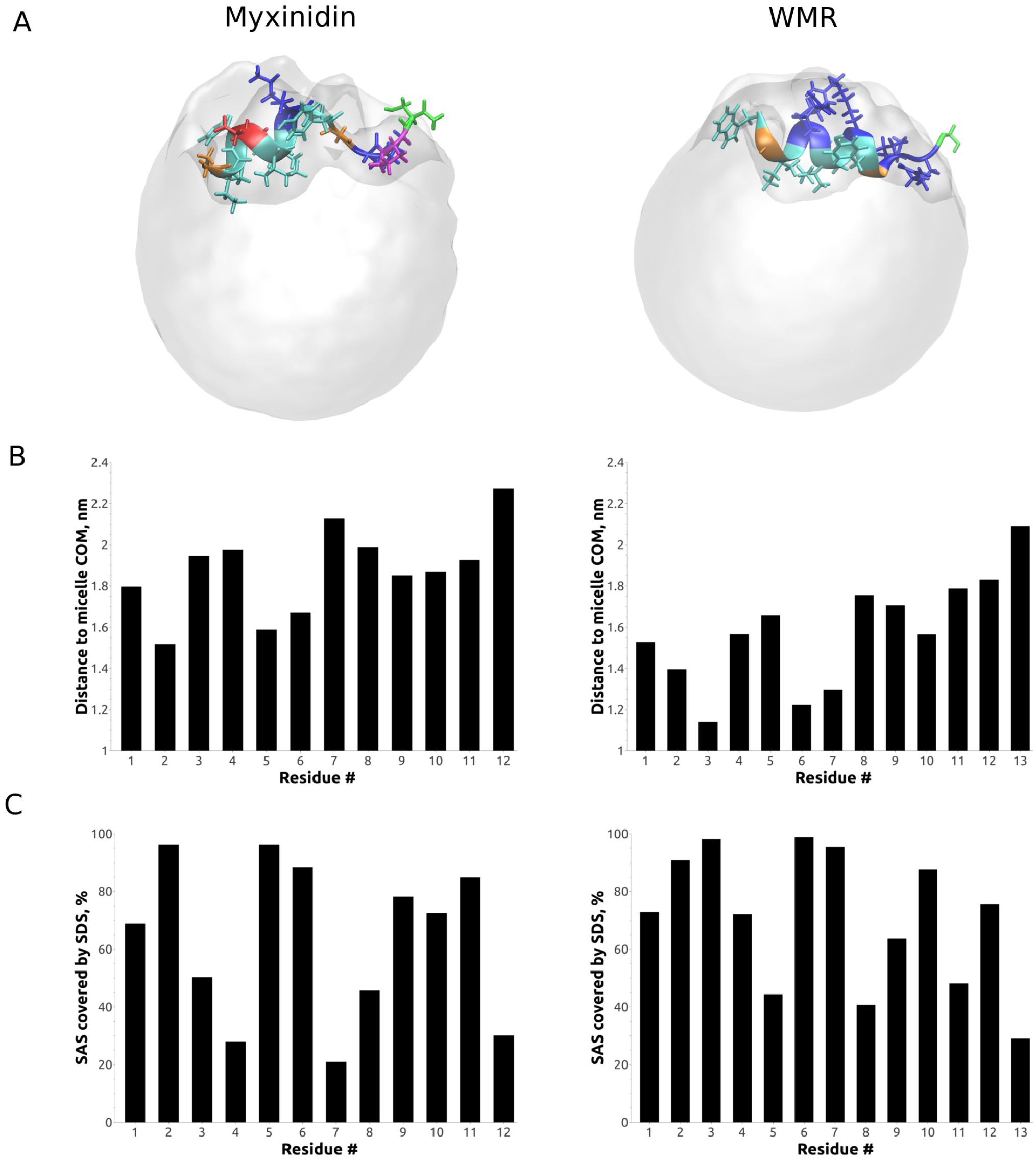
MD simulations of myxinidin and WMR in the presence of a SDS micelle. A) Ribbon representation of a representative structure (middle of top 1 cluster based on gromos clustering method with *gmx cluster* program) of Myxinidin (left) and WMR (right) in complex with the SDS micelle. The side chains are shown in stick representaiont. The color coding is the same as in Fig. 1A: cyan: aliphatic and aromatic, red: negatively charged, blue: positively charged, orange: glycine, magenta: proline, green: serine B) Analysis of the distance between the micelle COM and the residue COM for myxinidin (left) and WMR (right) as a function of the residue sequence position. C) Average solvent accessible surface (SAS) covered by SDS of myxinidin and WMR as a function of the residue sequence position.

Next, we calculated the distance between the center of mass of each residue and the micelle center of mass (Fig. 5B) for each peptide. All residues for both peptides reside mainly in the SDS micelle’s headgroup region, but WMR shows slightly deeper penetration into the micelle interior. Specifically, residues I2, I5, and L6 of myxinidin are located closer to the micelle COM, penetrating deeper into SDS’s hydrophobic core. At the same time, residues K7 and S12 are located closer to the micelle surface. Other residues of myxinidin reside at a similar distance between 1.8 to 2.0 nm from the micelle COM. A similar behavior is observed for WMR. Residues I3, I6, and L7 reside rather close to the SDS-micelle’s COM and hydrophobic core. Other residues exhibit very similar trends as observed with myxinidin, but reside a few angstroms closer to the micelle COM in absolute values, with S13 residue being the closest to the micelle surface.

Figure 5C shows the percentage of solvent accessible surface (SAS) of myxinidin or WMR covered by SDS molecules for different residues. As expected, residues immersed deeper into the SDS micelle interior show a higher percentage of SAS covered by SDS with a slight deviation from this trend with P11 residue of myxinidin and R12 residue of WMR. These residues lie closer to the micelle surface than the preceding K10 of myxinidin and K11 of WMR, but the SAS percentage covered by SDS is higher for these residues.

Overall, the residue distance to the micelle COM and the percentage of SAS covered by SDS molecules obtained from MD simulation support the data obtained with NMR PRE and chemical shift mapping that both peptides reside largely in the headgroup region of SDS micelle. A low percentage of SAS covered by SDS molecules for the S12 and D4 residues of myxinidin also agrees well with NMR PRE and chemical shift mapping data.

### MD simulations of multiple copies of myxinidin and WMR with DOPE/DOPG and DOPE/DOPG/CL membranes

To study the behavior of myxinidin and WMR in the presence of a negatively charged bilayer, we simulated these peptides with DOPE/DOPG and DOPE/DOPG/CL membranes for 5 microseconds. Eighteen copies of myxinidin or WMR were present in the simulation box to allow peptide-peptide interactions. Simulations were performed so that only one side of the membrane was accessible for the peptides, mimicking an initial stage of peptide-cell interaction when only the outer leaflet is exposed. As a control, we performed simulations of the same length of DOPE/DOPG and DOPE/DOPG/CL membranes without any peptides present.

Initially, we also performed our simulation in the presence of DMPC/DMPG membrane, but this bilayer composition turned out to be unstable at 303 K with the CHARMM36 force field. The DMPC/DMPG membrane exhibited a spontaneous transition from liquid to interdigitated gel phase after a few microseconds of the simulation, even without any peptides present. This transition happened faster for the system with myxinidin or WMR peptides (within the first 1-1.5 microseconds) compared to a pure membrane system (within ~4 microseconds), but there is not enough evidence to suggest that the peptides play a key role in this process. As a result, we do not show any data on DMPC/DMPG membrane setup.

Myxinidin and WMR peptides adopt a similar structure to the one observed in the micelle-associated state when bound to the membrane but tend to be less structurally stable when not in the membrane-bound state (Fig. 6).

**Fig. 6:**
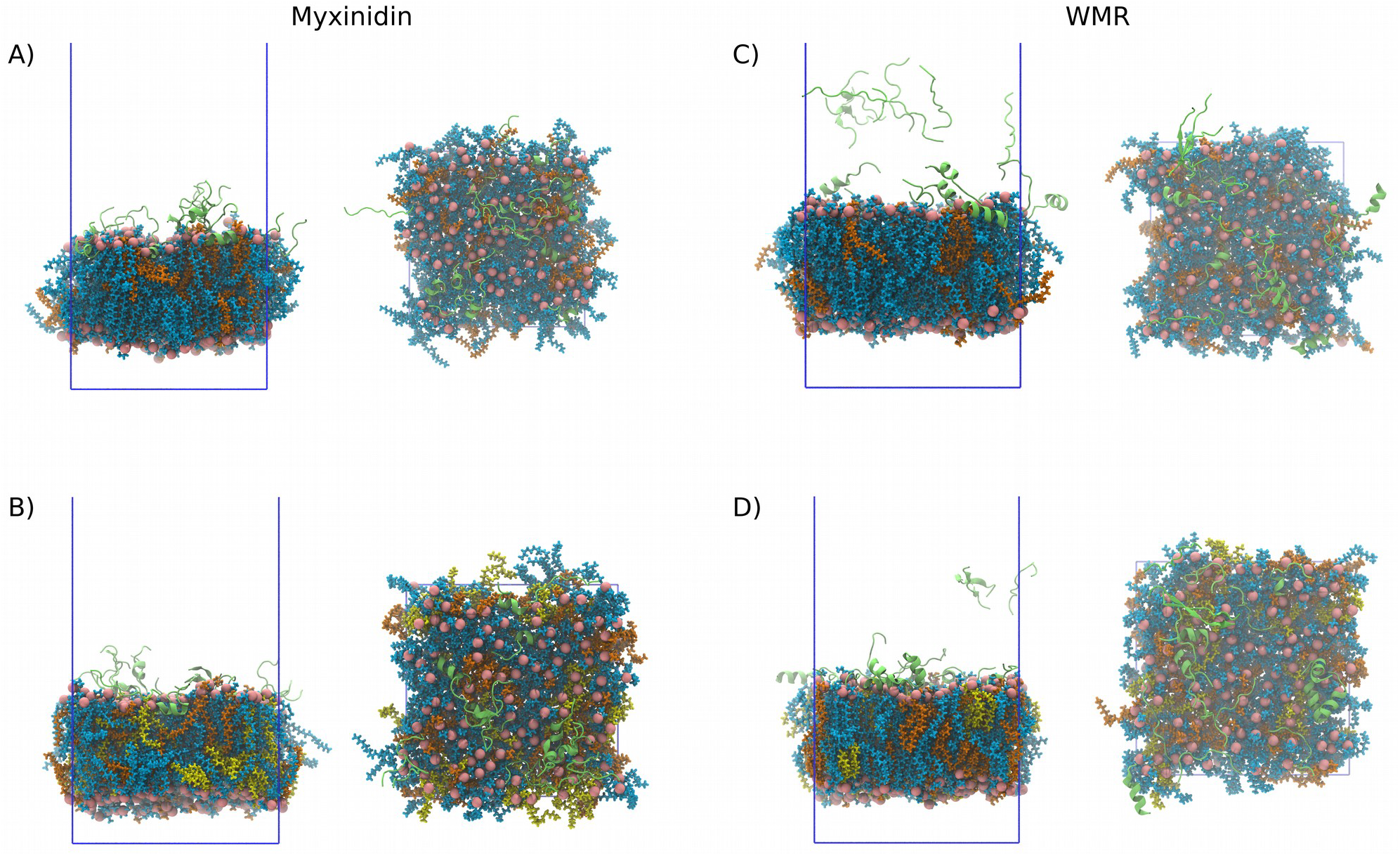
Snapshots for the MD simulations of multiple copies of myxinidin (A, B) and WMR (C, D) antimicrobial peptides in the presence of DOPE/DOPG (A, C) and DOPE/DOPG/CL (B, D) lipid bilayers. Peptides are shown in green. Different lipid types are represented in different colors (DOPE – blue, DOPG – orange, CL – yellow). Phosphate atoms of the lipid headgroups are shown with pink spheres. Side (left side of each sub figure) and top views (right side) of the simulated systems are shown. The periodic box is indicated by blue lines. Water and ions are not shown for clarity.

Figure 7 shows the density distribution of myxinidin and WMR peptides, together with DOPE/DOPG and DOPE/DOPG/CL membranes. Control data (bilayers simulated without any peptides, dashed lines in Fig. 7) are also shown for comparison. For both membrane compositions, membranebound peptides reside mainly in the lipid headgroup region, occasionally penetrating deeper towards the hydrophobic core of a membrane. Interestingly, the fraction of WMR that is bound to the membrane surface (first WMR density peak, Fig. 7 right two panels) is significantly affected by the membrane composition. This can be seen from the comparison of the magnitude of two WMR density peaks (turquoise lines, membrane-bound in the headgroup region and membrane-unbound about 5-8 nm from the bilayer center) in DOPE/DOPG and DOPE/DOPG/CL membrane cases (Fig. 7 A and B, right panels). Indicated by the increase in peak height around 2 nm, the presence of cardiolipin increases the fraction of WMR peptides that are bound to the lipid membrane.

**Fig. 7:**
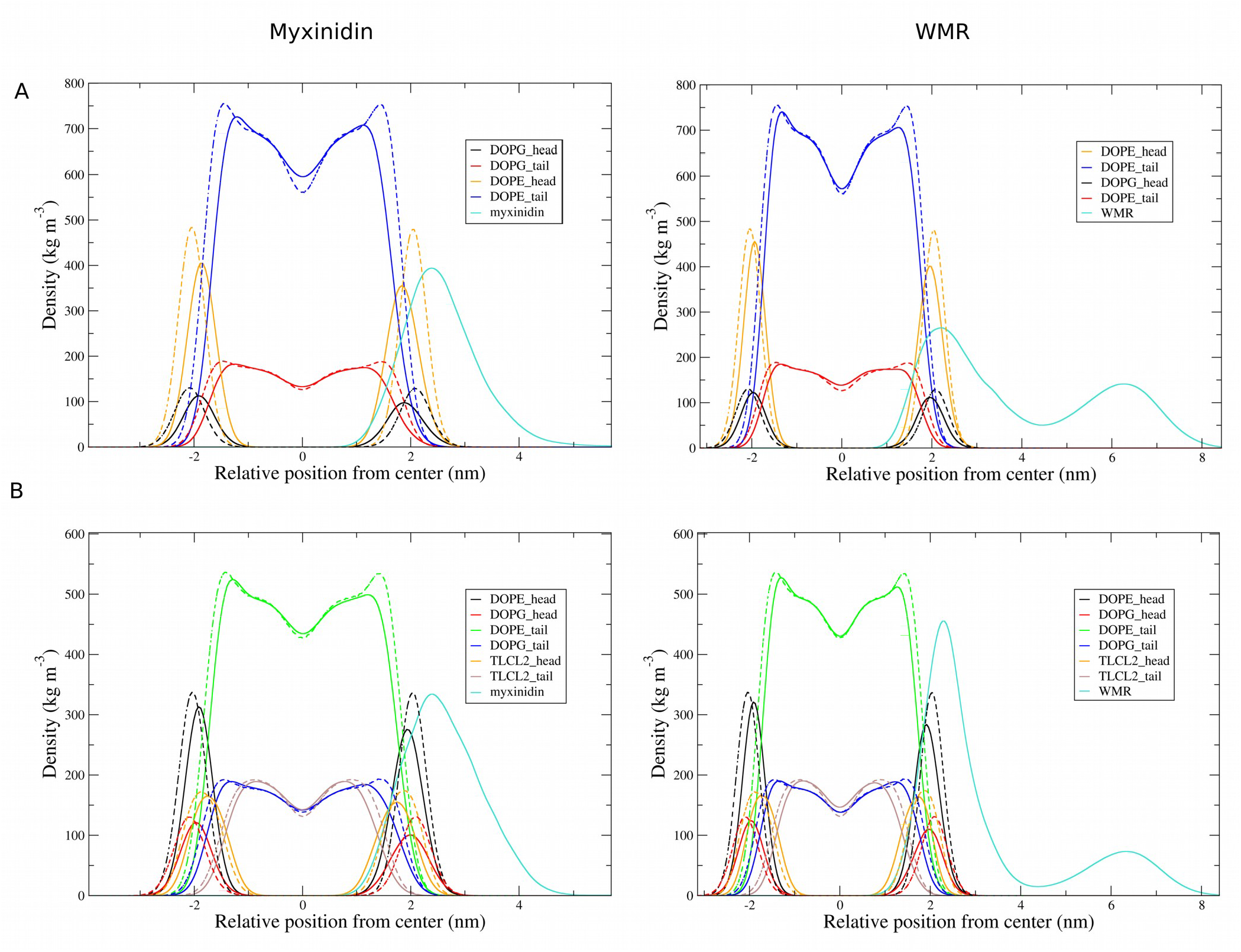
Density distribution of myxinidin and WMR peptides for the MD simulations in the presence of DOPE/DOPG (A) and DOPE/DOPG/CL (B) membranes. The color coding is described in the legend in each figure. The Control data (membrane simulated without any peptides) is shown as a dashed lines for comparison.

The presence of either myxinidin or WMR has an apparent effect on the total membrane density distribution, compared to the control (Fig. 7, continuous versus dashed lines). The density peaks for different membrane components are shifted closer to the bilayer center if peptides are present, even in the bottom monolayer, which is not directly exposed to the peptides. Also, asymmetry in the lipid density between different monolayers is introduced by the peptides. The effect on the monolayer directly exposed to the peptides is higher and is not restricted to the lipid headgroups, but significantly changes the shape of the distribution of the lipid tail densities. The character of the lipid density alterations induced by myxinidin and WMR is similar for both DOPE/DOPG and DOPE/DOPG/CL membranes and manifests itself in a decreased overall bilayer thickness, which translates into increased area per lipid.

Figure 8A shows the distance to the bilayer COM as a function of the residue sequence position for myxinidin (left panels) and WMR (right panels) when interacting with DOPE/DOPG or DOPE/DOPG/CL bilayers. Only the peptides that are in direct contact with the membrane are considered in this analysis. Similar to what we observed in the SDS micelle case, the N-terminal end of both peptides penetrates deeper into the membrane interior. For myxinidin in the DOPE/DOPG bilayer, residues I2, I5, and L6 are located closer to the bilayer COM than neighboring residues, but with DOPE/DOPG/CL membrane, the penetration depth of the residues 5 to 8 is almost the same, showing an alignment more parallel to the membrane surface of this part of the peptide. Indicated by smaller distances to the bilayer COM, the C-terminal half, especially residues 7-10 of myxinidin penetrate deeper into the membrane if cardiolipin (CL) is present. For WMR in DOPE/DOPG bilayers, residues 1-3, and 6 are closer to the bilayer COM than the neighboring residues, whereas in DOPE/DOPG/CL bilayers this is the case for residues 1, 2, and 5. The average distance to the bilayer COM is larger for the DOPE/DOPG membrane compared to DOPE/DOPG/CL. If we compare the distance to the bilayer COM for myxinidin and WMR, WMR shows 0.1-0.15 nm deeper penetration into the hydrophobic core of either DOPE/DOPG or DOPE/DOPG/CL membrane (Fig. 8A, 6).

**Fig. 8:**
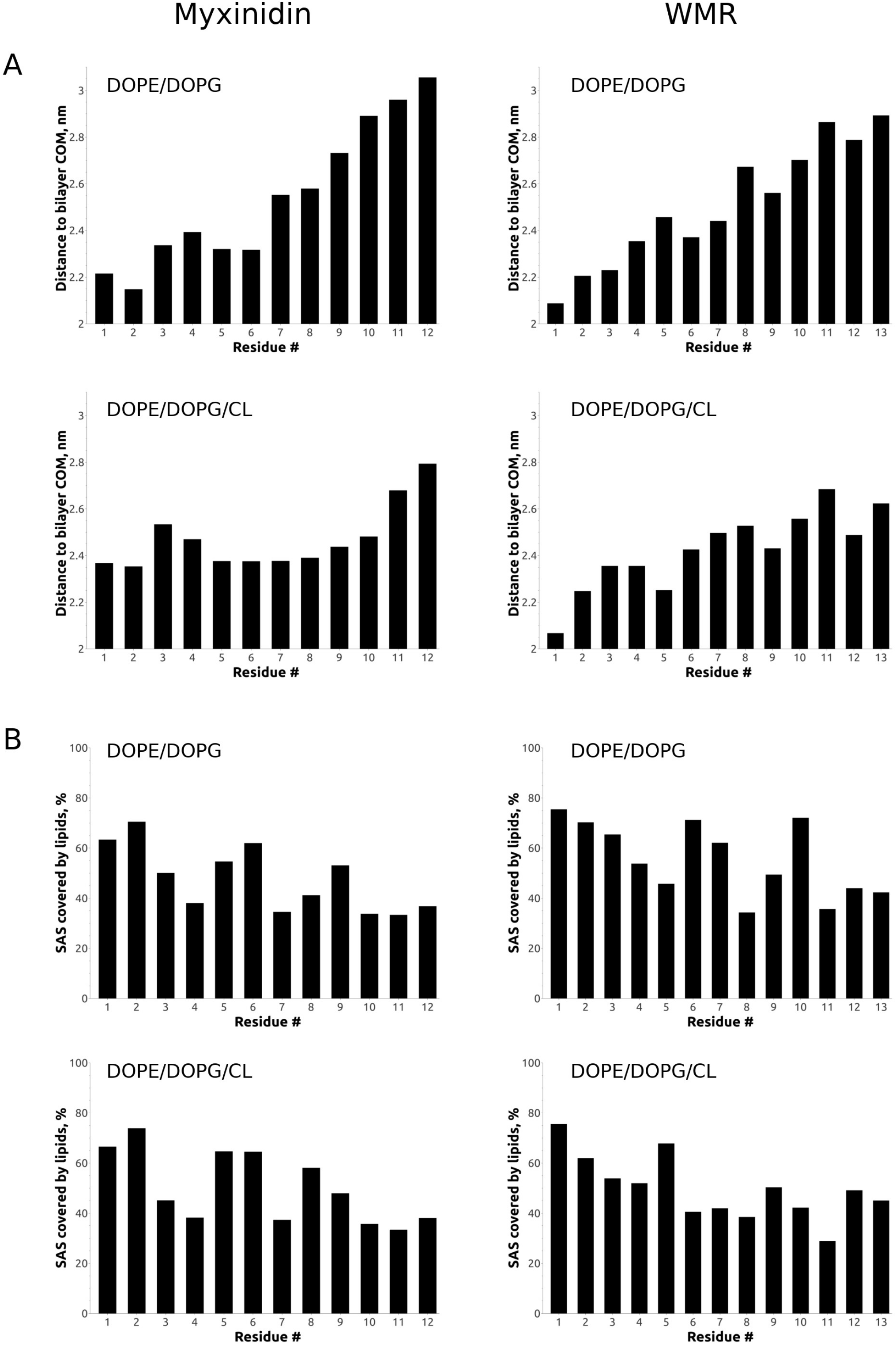
Information about the distance to the bilayer center the membrane covered areas for myxinidin and WMR from the MD simulations. A) Distance between the residue COM and bilayer center for myxinidin (left panel) and WMR (right panel) when interacting with DOPE/DOPG or DOPE/DOPG/CL membrane as a function of the residue sequence position. B) Average solvent accessible surface (SAS) covered by lipids of myxinidin and WMR as a function of the residue sequence position. Only peptides copies that are in direct contact with the membrane are considered in this analysis.

Figure 8B shows the SAS covered by lipids as a function of the residue sequence position. As with the penetration depth, only the peptides directly interacting with the bilayer are considered in the analysis. As expected, the residues that lie closer to the bilayer COM show a higher percentage of SAS covered by lipids. The exception to this trend occurs at the C-terminal end of both peptides – despite being located further away from the bilayer COM, the fraction of SAS covered by lipids is not much lower compared to neighboring residues. Also, residue K7 of myxinidin with DOPE/DOPG/CL membrane shows a lower percentage of SAS covered by lipids while being located at almost the same distance to bilayer COM as its neighbors.

In order to better understand the propensity of the peptides to self-interact and form clusters, we calculated the probability that a randomly chosen peptide belongs to a cluster of a size 1 (no aggregation) to 18 (all the peptide copies form a single aggregate) (Fig. 9). Two peptides were considered to belong to the same aggregate if they have contacts within 0.3 nm. For both myxinidin and WMR, the occurrence of self-interaction is high, but myxinidin has a significantly higher probability of forming large clusters. In contrast, WMR tends to form smaller clusters with both lipid membrane compositions. The majority of myxinidin copies belong to clusters of size 10 to 18, but WMR mainly forms smaller aggregates of a size below 10-11, especially with the DOPE/DOPG membrane. Interestingly, with DOPE/DOPG/CL membrane composition, both peptides show a tendency to form larger clusters compared to the DOPE/DOPG case (Fig. 9, 6).

**Fig. 9:**
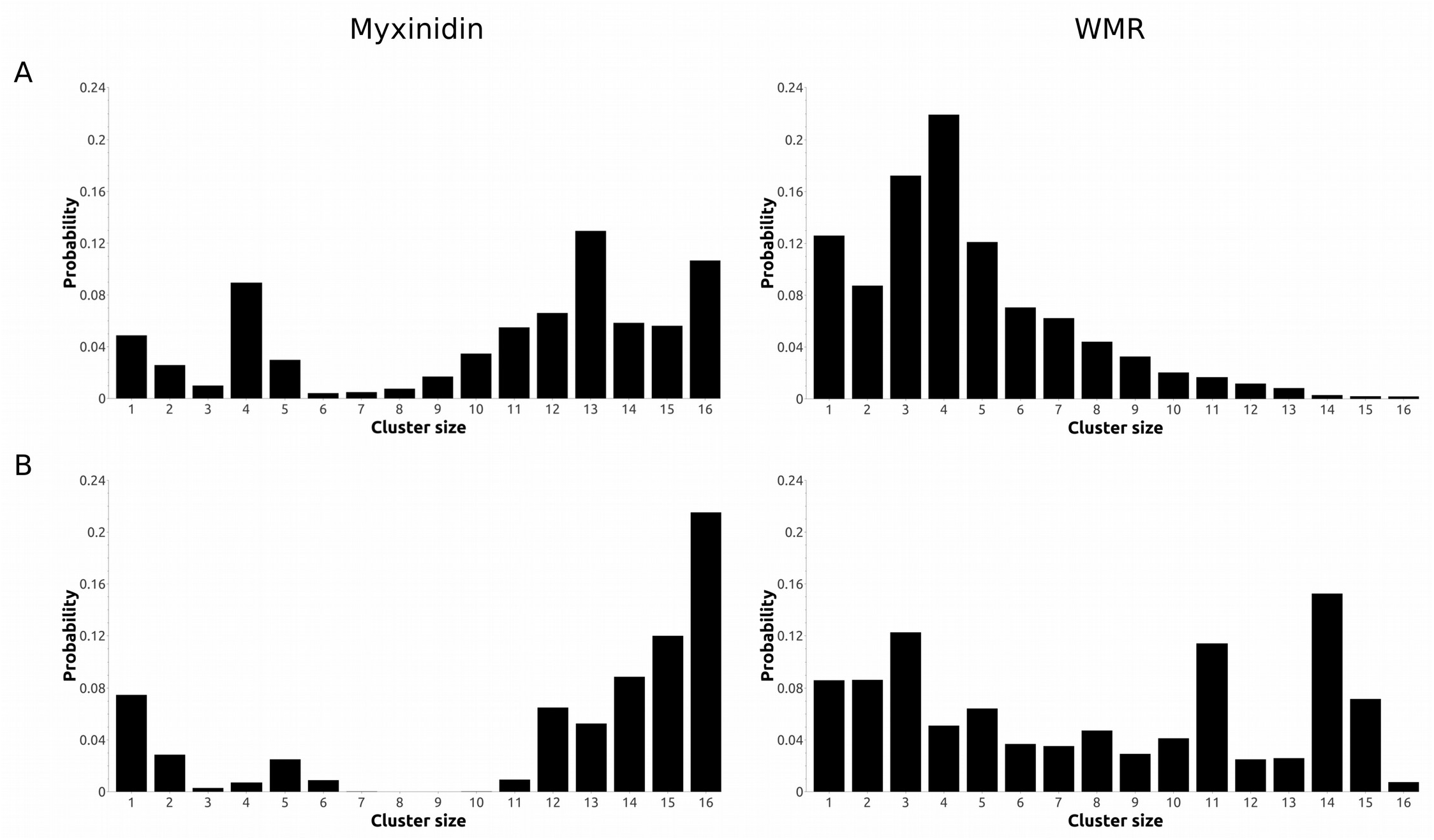
Analysis of the clustering propensity/probability from MD simulations of myxinidin and WMR antimicrobial peptides in the presence of DOPE/DOPG (A) and DOPE/DOPG/CL (B) membranes. Two peptides were considered to belong to the same aggregate if they have contacts within 0.3 nm. Cluster of a size 1 means there are no contacts of a given peptide copy within 0.3 nm with other peptides. Cluster of a size 18 means all the peptide copies belong to a single aggregate.

## Discussion

Both experimental data and simulations agree well that myxinidin and WMR adopt an α-helical structure in the presence of negatively charged membrane mimetics (Fig. 3, Fig. 5, SI Fig. S10). The importance of the α-helical structure-stabilizing effect of the negatively charged membrane mimetics is highlighted in the simulations with multiple copies of the peptides, where a certain fraction of the peptides are in the membrane-unbound state. We observe partial unfolding and structural instability of the peptides that are not in direct contact with the negatively charged headgroup region of DOPE/DOPG or DOPE/DOPG/CL membrane, but when the peptide is close to the membrane surface α-helical structure is restored (Fig. 6).

NMR PRE and chemical shift mapping data, together with simulations, indicate that the peptides reside mainly in the headgroup region of SDS or negatively charged bicelles/bilayers studied in this paper. Per residue distances to the micelle COM or bilayer COM together with SAS covered by SDS or bilayer lipids from the simulations show a good correlation with PRE and chemical shift mapping data (Fig. 4B, 5, and 8), with slight deviations for some residues. This deviation can be explained by the fact that these quantities, while being tightly related, are influenced by different factors. The experimental signal is highly dependent on the position of the nitroxide group of 5- and 16-SASL in a bilayer or SDS micelle. It is known that it resides near the lipid molecules’ headgroup region, so it is reasonable to assume that the peptide residue close to the nitroxide group of 5- or 16-SASL should be closer to the bilayer/micelle COM and have more SAS covered by lipids. Though it is not always a strict rule – in our simulations with SDS micelles, we observe an SDS molecule occasionally partially leaving the micelle and interacting with myxinidin or WMR residues that are solvent-exposed. This transient mode of interaction will give rise to SAS covered by SDS for the residues that lie further away from the micelle COM and the SDS headgroup region where the nitroxide group of 5- or 16-SASL is located.

The simulations of multiple copies of myxinidin and WMR with the DOPE/DOPG and DOPE/DOPG/CL membranes allowed us to observe collective modes of membrane-peptide and peptide-peptide interaction directly. All the copies of myxinidin were located near the membrane’s headgroup region during our simulations (Fig. 6, 7). Contrary, WMR shows two distinct groups of peptides – one near the membrane’s headgroup region, which corresponds to the membrane-bound state, and a second one in the water bulk. A higher positive charge carried by WMR can explain this observation – positively charged peptides disfavor close contacts with each other. The membrane’s negative charge partially counters this repulsive force, but with WMR, this balance is shifted compared to myxinidin. As a result, a higher fraction of WMR is located in the bulk water. This mechanism is further supported by the increase of the fraction of the WMR peptides located close to the membrane if we compare DOPE/DOPG and DOPE/DOPG/CL membranes (Fig. 6, 7). Both DOPE/DOPG and DOPE/DOPG/CL membranes are negatively charged, but an increased abundance of negative charge carried by cardiolipin promotes increased membrane affinity of WMR.

The difference between the modes of self-interaction of myxinidin and WMR is illustrated with the peptide self-aggregation data (Fig. 9). Myxinidin tends to form larger aggregates compared to WMR. This observation is in line with our speculation that WMR with the total charge of +6 is less prone to form close contacts with other copies of itself compared to myxinidin, which carries the total charge of +2. Interestingly, the presence of cardioplipin in DOPE/DOPG bilayers seems to promote close self-interactions and the formation of larger clusters for both peptides (Fig. 9). One of the reasons behind this observation could be that DOPE/DOPG/CL membrane composition carries higher negative charge density in the headgroup region compared to the DOPE/DOPG membrane composition, thus providing partial shielding of positive charges carried by the peptides thereby enabling close self-interaction.

Despite having multiple copies of WMR and myxinidin in our simulations with DOPE/DOPG and DOPE/DOPG/CL membranes and relatively long simulation trajectories (5 microseconds for each run), we failed to observe peptide-induced membrane disruption directly. As we mention in the Results section, we also performed DMPC/DMPG membrane simulations, but a pure DMPC/DMPG membrane turned out to spontaneously undergo a phase transition from liquid to interdigitated gel phase at 303K even without any peptides present. Though the transition happened significantly faster in the presence of the peptides, which can be a sign that peptides promote phase transition in DMPC/DMPG membranes, it can be just a coincidence and a simulation artifact. For DOPE/DOPG and DOPE/DOPG/CL membranes, we observe significant membrane thinning in the presence of myxinidin or WMR, which decrease the membrane stability. Despite the lack of direct observation of cooperative penetration deep into the membrane interior or pore formation by the peptides (Fig. 6), it is hard to rule out such a possibility if the simulations are extended for a longer period of time. At the same time, together with membrane thinning, this can be interpreted as a suggestion that myxinidin and WMR act via a carpet-like mechanism of membrane disruption. More extensive simulations, probably using enhanced sampling techniques like replica-exchange, are required to answer this question fully.

The modes of interaction observed in the current study can also be related to myxinidin and WMR ability to disrupt DOPE/DOPG and DOPE/DOPG/CL membranes measured in Lombardi *et al.^22^*. At the peptide/lipid ratio of 1 to 10, as in our simulations, myxinidin shows significantly lower leakage of the fluorophores from the model vesicles with DOPE/DOPG/CL membrane composition compared to WMR. On the contrary, both peptides show a similar percentage of the fluorophore leakage with DOPE/DOPG membrane composition. This difference in the ability to disrupt membranes of different composition can be related to a different effective concentration of the peptides in the bilayer’s headgroup region. Myxinidin density in the headgroup region of the DOPE/DOPG/CL membrane is lower compared to the DOPE/DOPG membrane (Fig. 6, 7). Also, myxinidin tends to form larger aggregates with DOPE/DOPG/CL membrane composition (Fig. 9). These observations can indicate that myxinidin, when exposed to the DOPE/DOPG/CL membrane, forms a smaller number of close contacts with the membrane compared to the DOPE/DOPG membrane. Instead, more peptides are in the big aggregates that directly interact with the membrane only with its edges – lots of the peptides are trapped inside an aggregate. Thus, a smaller fraction of the membrane surface is covered by myxinidin, which is also supported by smaller myxinidin density in the headgroup region of the DOPE/DOPG/CL membrane compared to DOPE/DOPG membrane composition. If myxinidin acts through the carpet-like mechanism of membrane disruption, this would lead to the lower leakage of the fluorophores from the model vesicles.

## Conclusions

We determined the structures of the antimicrobial peptides myxinidin and WMR associated with bacterial membrane mimetic micelles and bicelles by NMR, CD spectroscopy, and Molecular Dynamics simulations. Both peptides were found to have a mostly α-helical structure in the presence of negative membrane mimetics. Myxinidin and WMR reside mainly in the headgroup region of the membrane or SDS micelle and have a noticeable membrane thinning effect on the overall bilayer structure. Myxinidin and WMR show a different tendency to self-aggregate that depends on the membrane composition, which may be related to the previously observed difference in the peptides’ ability to disrupt different types of model membranes and to the different antimicrobial activity observed for different types of gram-positive and gram-negative bacteria^19–21^.

## Supporting information

Top 1 pdb structure of myxinidin in SDS

Ensemble of best 20 structures of myxinidin in SDS

Ensemble of best 20 structures of myxinidin in bicelles

Top 1 pdb structure of WMR in SDS

Ensemble of best 20 structures of WMR in SDS

Ensemble of best 20 structures of WMR in bicelles

## Abbreviations

CL: cardiolipin
CMC: critical micelle concentration
DihepPC: 1,2-diheptanoyl-*sn*-glycero-3-phosphocholine
DMPC: 1,2-dimyristoyl-*sn*-glycero-3-phosphocholine
DMPG: 1,2-dimyristoyl-*sn*-glycero-3-phospho-(1’-rac-glycerol)
DOPE: 1,2-dioleoyl-sn-glycero-3-phosphoethanolamine
DOPG: 1,2-dioleoyl-sn-glycero-3-phospho-(1’-rac-glycerol)
SDS: sodium dodecyl sulfate
SI: supplementary information
MD: molecular dynamics
CD: circular dichroism
NMR: nuclear magnetic resonance
PRE: paramagnetic relaxation enhancement

## Acknowledgment

S.A.D. is grateful to Prof. Dr. M. Sattler and Prof. Dr. B. Reif from the Technische Universität München/Helmholtz Zentrum München for hosting her group and for sharing their facilities.

## Supporting Information

CD spectra of myxinidin and WMR in SDS micelles and DMPC/DMPG/CL bicelles. Plots of the amide region of the 2D ^1^H-^1^H TOCSY and/or NOESY spectra of myxinidin and WMR in SDS micelles and DMPC/DMPG/CL bicelles labeled with the corresponding assignments and a table listing all assigned ^1^H chemical shifts. Plots of the superpositions of the amide region of the 2D ^1^H-^1^H TOCSY of myxinidin and WMR in negatively charged SDS micelles and DMPC/DMPG/CL bicelles in the absence and presence of spin-labeled stearic acid (5- or 16-SASL). Plots of myxinidin and WMR secondary structure as a function of simulation time. Snapshots of the simulations of multiple copies of WMR and myxinidin interacting with a DOPE/DOPG/CL bilayer.

