## Supplemental information for "Structural characterization of the antimicrobial peptides myxinidin and WMR in bacterial membrane mimetic micelles and bicelles"

### Supplementary Information

#### Supplementary Tables

**Table S1** Chemical shift assignment of myxinidin and WMR in negatively charged membrane mimetic micelles and bicelles

| Myxinidin G1-I2-H3-D4-I5-L6-K7-Y8-G9-K10-P12-S13 | | WMR W1-G2-I3-R4-R5-I6-L7-K8-Y9-G10-K11-R12-S13 | |
|---|---|---|---|
| 150 mM SDS | DMPG/DMPC/CL/DihepPC bicelles | 150 mM SDS | DMPG/DMPC/CL/DihepPC bicelles |
| | | 1.HA 4.445 1 | 1.HA 4.442 1 |
| | | 1.HB2 3.441 2 | 1.HB2 3.280 2 |
| | | 1.HB1 3.556 2 | 1.HB1 3.380 2 |
| | | 1.HD1 7.428 1 | 1.HD1 7.440 1 |
| | | 1.HE1 10.049 2 | 1.HE1 10.520 2 |
| | | 1.HZ2 7.480 2 | 1.HZ2 7.480 2 |
| | | 1.HH2 7.145 1 | 1.HH2 7.085 1 |
| | | 1.HE3 7.685 2 | 1.HZ3 6.982 2 |
| | | 1.HZ3 7.080 2 | 1.HE3 7.550 2 |
| 1.HA2 4.060 2 | 1.HA1 3.972 2 | 2.HA1 4.074 2 | 2.HA1 4.053 2 |
| 1.HA1 4.193 2 | | | |
| 2.HN 8.831 1 | 2.HN 9.026 1 | 3.HN 8.046 1 | 3.HN 8.433 1 |
| 2.HA 3.984 1 | 2.HA 4.062 1 | 3.HA 3.991 1 | 3.HA 4.088 1 |
| 2.HB 1.932 1 | 2.HB 1.879 1 | 3.HB 2.044 1 | 3.HB 1.935 1 |
| 2.HG12 1.316 9 | 2.HG12 1.270 1 | 3.HG12 1.612 9 | 3.HG21 0.980 4 |
| 2.HG11 1.539 9 | 2.HG11 1.370 1 | 3.HG11 1.695 9 | |
| 2.MD1 0.922 1 | 2.MD1 0.862 1 | 3.MD1 0.945 1 | |
| 2.MG2 0.965 4 | 2.MG2 0.901 2 | 3.MG2 0.972 4 | |
| 3.HN 8.660 1 | 3.HN 8.765 1 | 4.HN 8.144 1 | 4.HN 8.585 1 |
| 3.HA 4.371 1 | 3.HA 4.533 1 | 4.HA 4.060 1 | 4.HA 4.055 1 |
| 3.HB1 3.310 2 | 3.HB1 3.262 2 | 4.HB2 1.867 2 | 4.HB1 1.924 2 |
| 3.HD2 7.386 2 | 3.HD2 7.278 2 | 4.HB1 1.947 2 | 4.HG1 1.771 2 |
| 3.HE1 8.595 2 | 3.HE1 8.404 2 | 4.HG1 1.663 2 | |
| | | 4.HD1 3.219 2 | |
| | | 4.HE 7.177 1 | |
| 4.HN 7.811 1 | 4.HN 7.941 1 | 5.HN 7.845 1 | 5.HN 8.071 1 |
| 4.HA 4.493 1 | 4.HA 4.498 1 | 5.HA 4.135 1 | 5.HA 4.157 1 |
| 4.HB1 2.860 2 | 4.HB1 2.767 2 | 5.HB1 1.941 2 | 5.HB1 1.925 2 |
| | | 5.HG2 1.708 2 | 5.HG1 1.805 2 |
| | | 5.HG1 1.741 2 | |
| | | 5.HD1 3.226 2 | |
| | | 5.HE 7.180 1 | |

| | | | |
|---|---|---|---|
| 5.HN 7.578 1 | 5.HN 7.830 1 | 6.HN 7.841 1 | 6.HN 7.796 1 |
| 5.HA 4.020 1 | 5.HA 4.065 1 | 6.HA 3.992 1 | 6.HA 4.054 1 |
| 5.HB 2.028 1 | 5.HB 1.980 1 | 6.HB 2.052 1 | 6.HB 2.032 1 |
| 5.HG12 1.336 9 | 5.HG12 1.328 1 | 6.HG12 1.330 9 | 6.HG11 1.705 9 |
| 5.HG11 1.696 9 | 5.HG11 1.614 9 | 6.HG11 1.697 9 | 6.MG2 0.977 4 |
| 5.MD1 0.919 1 | 5.MD1 0.906 1 | 6.MD1 0.940 1 | |
| 5.MG2 0.993 4 | 5.MG2 0.957 4 | 6.MG2 1.013 4 | |
| 6.HN 7.766 1 | 6.HN 7.971 1 | 7.HN 7.846 1 | 7.HN 7.874 1 |
| 6.HA 4.260 1 | 6.HA 4.287 1 | 7.HA 4.186 1 | 7.HA 4.195 1 |
| 6.HB1 1.812 2 | 6.HB1 1.745 2 | 7.HB1 1.860 2 | 7.HB1 1.824 2 |
| 6.HG 1.636 1 | 6.HG 1.595 1 | 7.HB2 1.547 2 | 7.MD1 0.990 1 |
| 6.MD1 0.931 4 | 6.HD11 0.951 2 | 7.HG 1.590 1 | |
| 6.MD2 0.961 4 | 6.HD21 0.877 2 | 7.MD1 0.970 4 | |
| | | 7.MD2 0.940 4 | |
| 7.HN 7.699 1 | 7.HN 7.821 1 | 8.HN 7.808 1 | 8.HN 7.792 1 |
| 7.HA 4.133 1 | 7.HA 4.211 1 | 8.HA 4.081 1 | 8.HA 4.183 1 |
| 7.HB1 1.634 2 | 7.HB1 1.702 2 | 8.HB1 1.746 2 | 8.HB1 1.925 2 |
| 7.HG1 1.156 2 | 7.HG1 1.248 2 | 8.HG2 1.115 2 | 8.HD1 1.625 2 |
| 7.HD1 1.604 2 | 7.HD1 1.630 2 | 8.HG1 1.318 2 | 8.HE1 2.971 2 |
| 7.HE1 2.940 2 | 7.HE1 2.966 2 | 8.HD1 1.617 2 | |
| | | 8.HE1 2.955 2 | |
| 8.HN 7.829 1 | 8.HN 7.944 1 | 9.HN 7.858 1 | 9.HN 7.882 1 |
| 8.HA 4.565 1 | 8.HA 4.543 1 | 9.HA 4.417 1 | 9.HA 4.474 1 |
| 8.HB2 3.010 2 | 8.HB2 2.986 2 | 9.HB2 3.062 2 | 9.HB2 3.031 2 |
| 8.HB1 3.224 2 | 8.HB1 3.137 2 | 9.HB1 3.221 2 | 9.HB1 3.134 2 |
| 8.HD1 7.239 2 | 8.HD1 7.175 2 | 9.HD1 7.191 2 | 9.HD1 7.174 2 |
| 8.HE1 6.866 2 | 8.HE1 6.852 2 | 9.HE1 6.843 2 | 9.HE1 6.832 2 |
| 9.HN 7.989 1 | 9.HN 8.236 1 | 10.HN 8.232 1 | 10.HN 8.313 1 |
| 9.HA1 3.983 2 | 9.HA1 3.912 2 | 10.HA1 3.955 2 | 10.HA1 3.902 2 |
| 10.HN 7.957 1 | 10.HN 8.045 1 | 11.HN 8.019 1 | 11.HN 8.066 1 |
| 10.HA 4.529 1 | 10.HA 4.643 1 | 11.HA 4.292 1 | 11.HA 4.046 1 |
| 10.HB1 1.887 2 | 10.HB1 1.869 2 | 11.HB2 1.861 2 | 11.HB1 1.920 2 |
| 10.HB2 1.741 2 | 10.HG1 1.488 2 | 11.HB1 1.941 2 | 11.HG1 1.530 2 |
| 10.HG2 1.521 2 | 10.HD1 1.748 2 | 11.HG1 1.526 2 | 11.HG2 1.482 2 |
| 10.HG1 1.544 2 | 10.HE1 3.036 2 | 11.HD1 1.735 2 | 11.HD1 1.721 2 |
| 10.HD2 1.772 2 | | 11.HE1 3.055 2 | 11.HE1 3.035 2 |
| 10.HD1 1.810 2 | | | |
| 10.HE1 3.064 2 | | | |
| 11.HA 4.519 1 | 11.HA 4.499 1 | 12.HN 8.145 1 | 12.HN 8.590 1 |
| 11.HB2 2.056 2 | 11.HB2 1.996 2 | 12.HA 4.296 1 | 12.HA 4.053 1 |
| 11.HB1 2.394 2 | 11.HB1 2.354 2 | 12.HB2 1.888 2 | 12.HB1 1.928 2 |
| 11.HG1 2.123 2 | 11.HG1 2.067 2 | 12.HB1 1.983 2 | 12.HG1 1.774 2 |
| 11.HD2 3.749 2 | 11.HD2 3.707 2 | 12.HG1 1.712 2 | |
| 11.HD1 3.905 2 | 11.HD1 3.869 2 | 12.HD1 3.230 2 | |
| 12.HN 8.273 1 | 12.HN 8.416 1 | 13.HN 8.030 1 | 13.HN 8.055 1 |
| 12.HA 4.420 1 | 12.HA 4.404 1 | 13.HA 4.360 1 | 13.HA 4.460 1 |
| 12.HB2 3.914 2 | 12.HB1 3.909 2 | 13.HB1 3.875 2 | 13.HB1 3.908 2 |
| 12.HB1 3.950 2 | 12.H2 7.194 2 | | |
| 12.H2 7.170 2 | 12.H1 7.620 2 | | |
| 12.H1 7.514 2 | | | |

#### Supplementary results

**Analysis of interatomic distances in the calculated NMR structures to estimate helix-stabilization by ionic or cation-π interations**

1) Myxinidin in the presence of negatively charged SDS micelles

Due to the i, i+3 spacing of G1 and K7 to D4 within the α-helix, salt bridge interactions may stabilize it. The side chain carboxyl oxygen atoms of D4 show a distance below 4 Å, a range typical for salt bridge interactions, to the amide protons of the N-terminal G1 in 9 and to that of K7 in 2 of the 20 lowest energy structures. However, since the NOE data do not directly restrain these potential interactions, the first option may occur more often. If the distance limit is increased to 5 Å, the side chain carboxyl oxygen atoms of D4 come close to the amide protons of the N-terminal G1 in 14 and K7 in 7 of the 20 lowest energy structures. The helical structure may further be stabilized by cation-π interactions between the aromatic rings of H3 and Y8 and the positively charged side chains of K7 or K10. In the 20 lowest energy structures, the carbon and/or nitrogen atoms of the aromatic ring of H3 show a distance lower than 6 Å, which is in the range typical for these interactions to occur, to the side-chain nitrogen of K 7 in 4 of 20 and that of Y8 to the side chain nitrogen atoms of K7 in 2 of 20 and of K10 in 0 of 20.

2) Myxinidin in the presence of negatively charged bicelles

Based on the analysis of the potential helix stabilizing effect from salt bridge interactions of the D4 side chain carboxyl oxygen atoms with amide protons of the N-terminal G1 or the side chain of K7, distances below 4 Å are found in 6 and 4 of the 20 lowest energy structures, respectively. Regarding a potential stabilization of the helix by cation-π interactions, distances below 6 Å between the heavy atoms of the aromatic ring of H3 to the side-chain nitrogen of K7 are found in 10 out of 20, between that of Y8 to the side chains nitrogens of K7 in 2 out of 20, and to K10 in 1 out of 20 lowest energy structures.

3) WMR in presence of negatively charged SDS micelles

Stabilization of the helical region might for WMR only occur by cation-π interactions between the present aromatic side chains (W1, Y9) and the positively charged ones (R4, R5, K8, K11, R12). In the micelle structure, distances below 6 Å between the heavy atoms of the aromatic ring of W1 to the side-chain terminal carbon and nitrogen atoms of R4 and R5 are found in 0 and 4 out of 20 lowest energy structures, respectively, and between that of Y9 and the side chains nitrogens of K8 in 6 and of K11 in 0 out of 20 and to the side-chain terminal carbon and nitrogen atoms of R5 in 5 and of R12 in 10 out of 20 lowest energy structures. In the bicelle structure, distances below 6 Å between the heavy atoms of the aromatic ring of W1 to the side-chain terminal carbon and nitrogen atoms of R4 and R5 are found in 0 and 10 of the 20 lowest energy structures, respectively, and between that of Y9 and the side chain nitrogens of K8 in 2 and of K11 in 1 of 20 and to the side-chain terminal carbon and nitrogen atoms of R5 in 5 and of R12 in 4 out of 20 lowest energy structures.

#### Supplementary figures

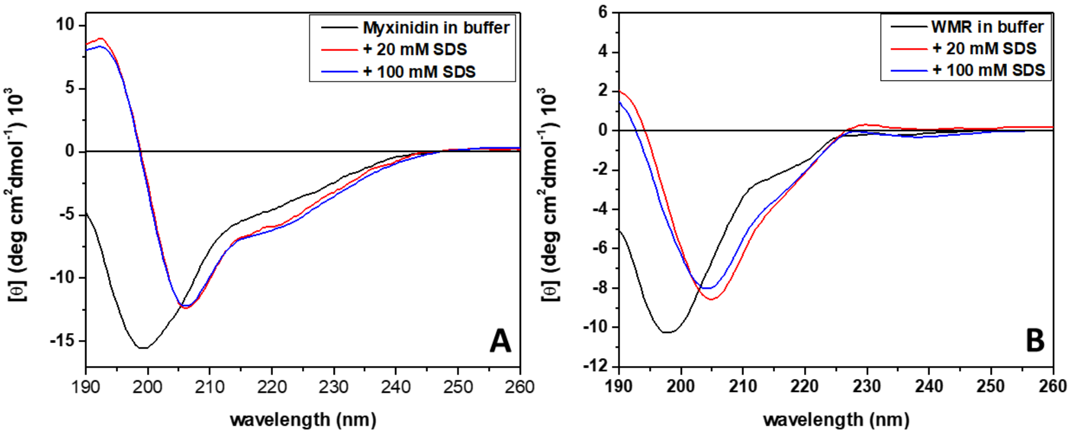


**Fig. S1:** Far-UV CD spectra for the peptides myxinidin (A) and WMR (B) in the absence (black lines) and in presence of 20 mM SDS (red lines) and 100 mM SDS (blue lines). All the spectra were recorded in 10 mM phosphate buffer, pH 7.4, at the temperature of 25 °C.

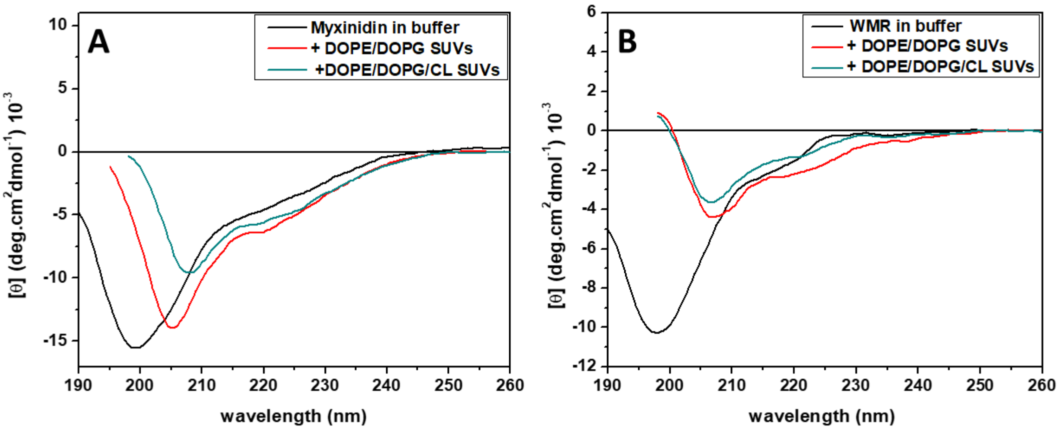


**Fig. S2:** Plots of the amide and aromatic region of the $^1$H-$^1$H 2D TOCSY (mixing time 70 ms) and NOESY (mixing time 100 ms) of myxinidin in negatively charged SDS micelles. The assigned amide and aromatic protons are labeled with the one-letter amino acid code, the residue sequence position and the atom name. The color coding is the same as in the amino acid sequence plot shown in Fig. 1A and in the structures in Fig. 2. The cross peaks belonging to a certain amide or aromatic proton are connected by vertical lines in the same color as the label. The $^1H$-$^1H$ 2D NOESY with a mixing time of 200 ms looked overall very similar and did not show more cross peaks, thus the 100 ms one was picked.

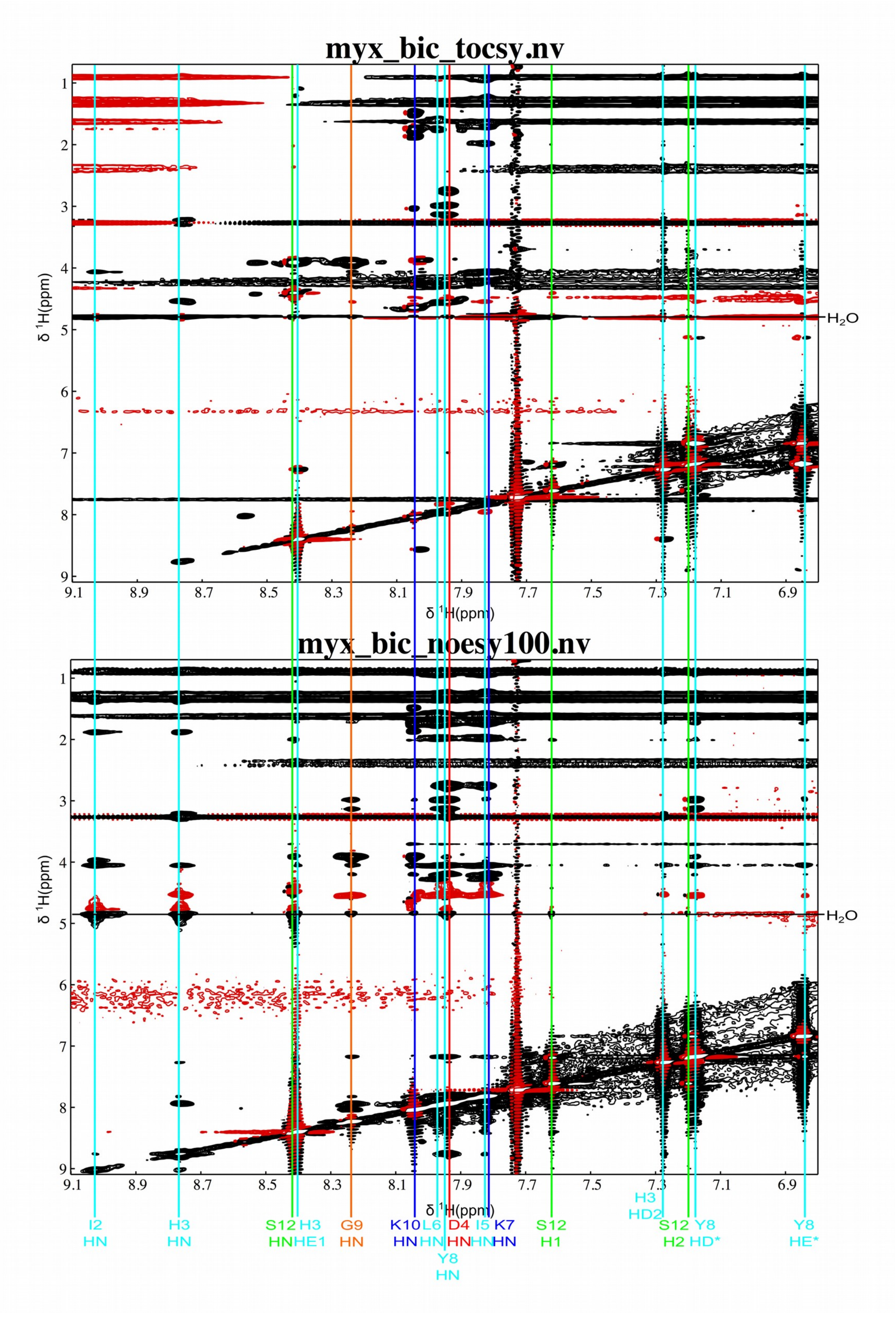

**Fig. S3:** Plots of the amide and aromatic region of the $^{1}$H-$^{1}$H 2D TOCSY (mixing time 70 ms) and NOESY (mixing time 100 ms) of myxinidin in negatively charged DMPC/DMPG/CL bicelles. The assigned amide and aromatic protons are labeled with the one-letter amino acid code, the residue sequence position and the atom name. The color coding is the same as in the amino acid sequence plot shown in Fig. 1A and in the structures in Fig. 2. The cross peaks belonging to a certain amide or aromatic proton are connected by vertical lines in the same color as the label. The $^{1}$H-$^{1}$H 2D NOESY with a mixing time of 200 ms looked overall very similar and did not show significantly more cross peaks, thus the 100 ms one was picked. Note that for the preparation of bicelles only deuterated DMPC ($d_{54}$) was used, whereas DMPG, CL and DihepPC were fully protonated. In case of SDS micelles, only deuterated SDS ($d_{25}$) was used. Thus, the spectra in SDS micelles (Fig. S2) look better overall.

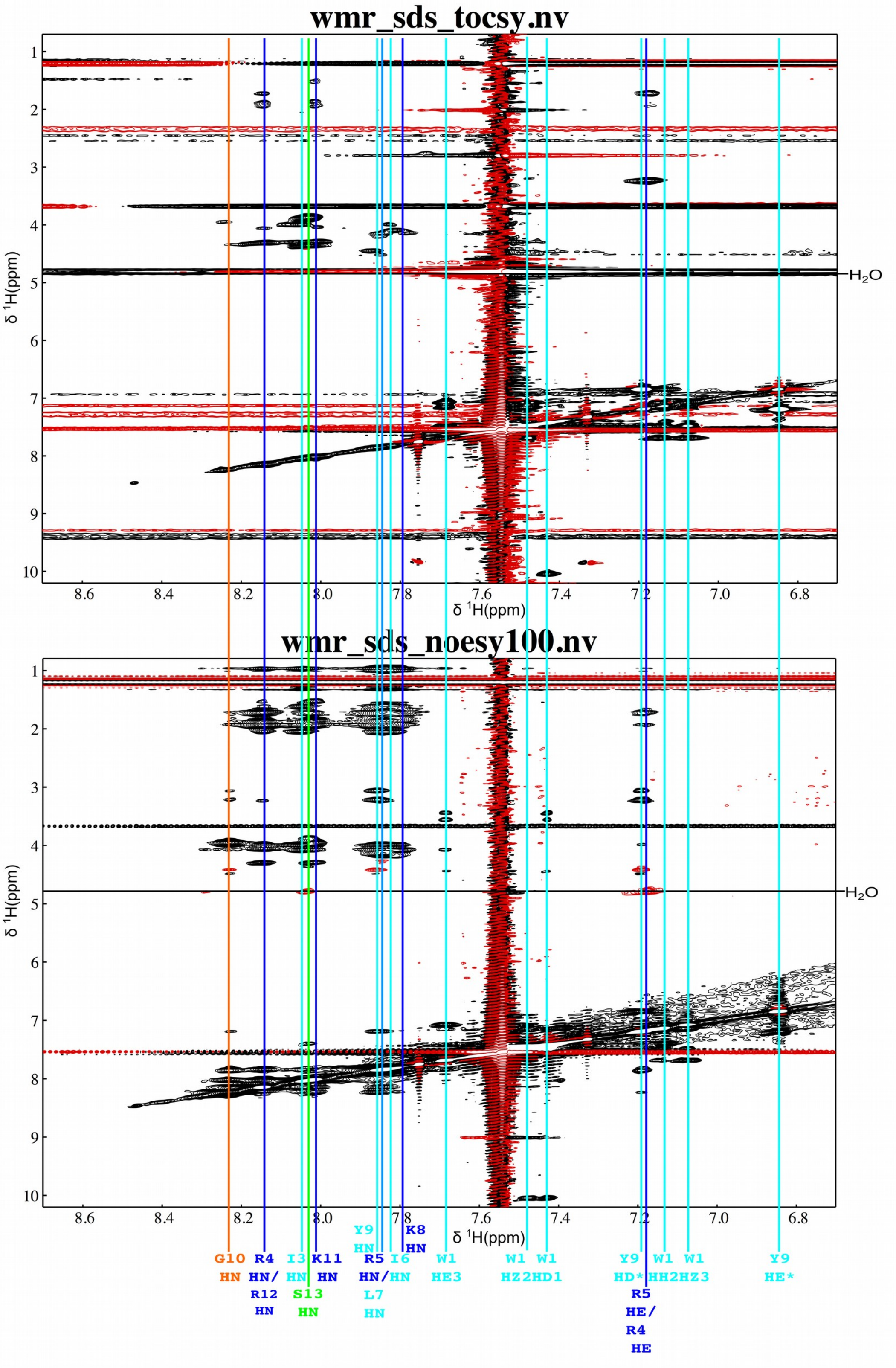

wmr_sds_tocsy.nv
wmr_sds_noesy100.nv
δ 1H(ppm)
δ 1H(ppm)
H2O
G10 HN
R4 HN/ R12 HN
I3 HN
S13 HN
K11 HN
Y9 HN
R5 HN/ L7 HN
I6 HN
K8 HN
W1 HE3
W1 HZ2
W1 HD1
Y9 HD*
R5 HE/ R4 HE
W1 HH2
W1 HZ3
Y9 HE*

**Fig. S4:** Plots of the amide and aromatic region of the $^{1}$H-$^{1}$H 2D TOCSY (mixing time 70 ms) and NOESY (mixing time 100 ms) of WMR in negatively charged SDS micelles. The assigned amide and aromatic protons are labeled with the one-letter amino acid code, the residue sequence position and the atom name. The color coding is the same as in the amino acid sequence plot shown in Fig. 1A and in the structures in Fig. 2. The cross peaks belonging to a certain amide or aromatic proton are connected by vertical lines in the same color as the label. The $^{1}$H-$^{1}$H 2D NOESY with a mixing time of 200 ms looked overall very similar and did not show more cross peaks; however, they exhibited more signal ridges and distortions. Thus the 100 ms spectrum was used.

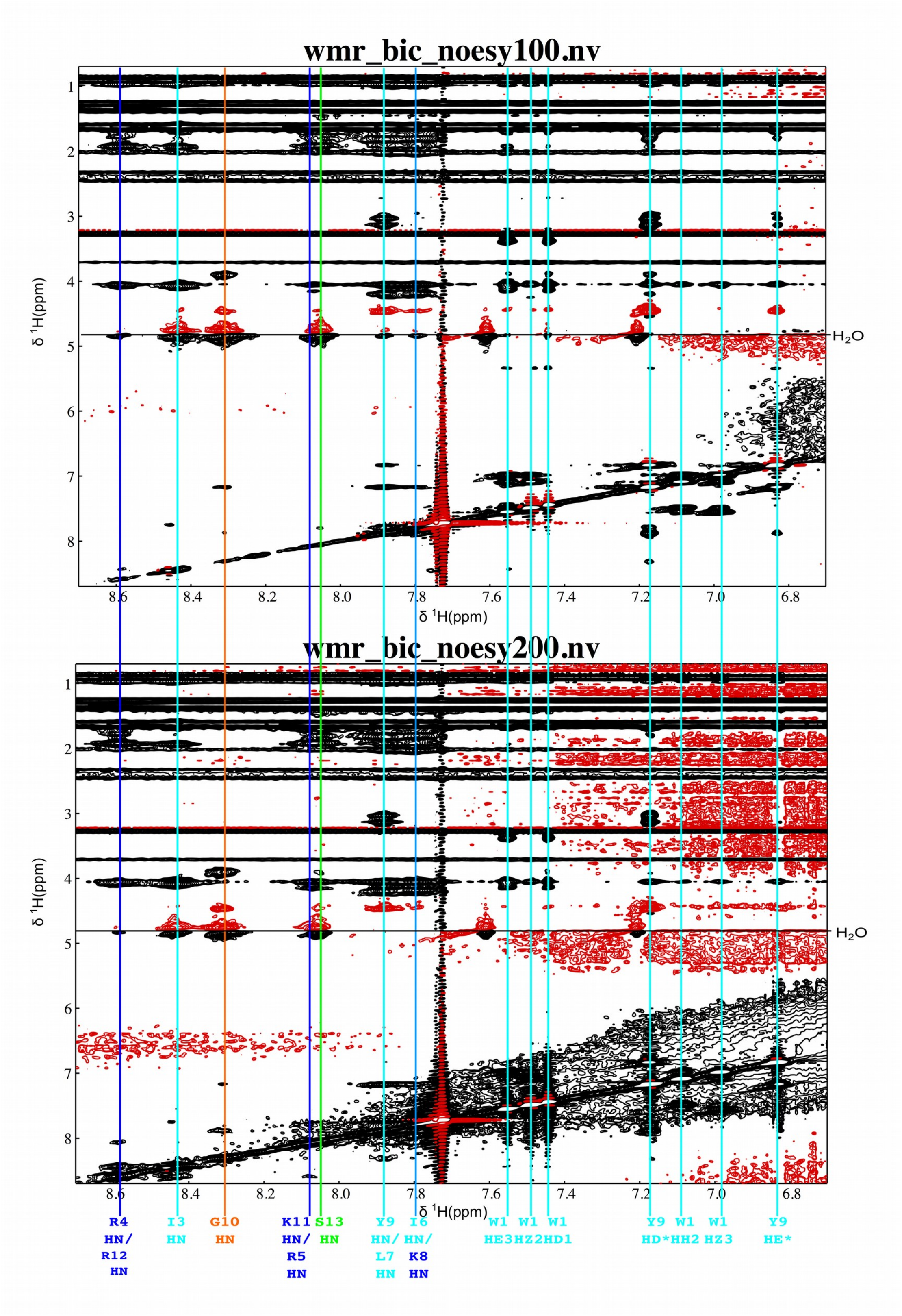


**Fig. S5:** Plots of the amide and aromatic region of the $^{1}H$-$^{1}H$ 2D NOESY spectra (mixing time 100 ms top and 200 ms bottom) of WMR in negatively charged DMPC/DMPC/CL bicelles. The assigned amide and aromatic protons are labeled with the one-letter amino acid code, the residue sequence position and the atom name. The color coding is the same as in the amino acid sequence plot shown in Fig. 1A and in the structures in Fig. 2. The cross peaks belonging to a certain amide or aromatic proton are connected by vertical lines in the same color as the label. The $^{1}H$-$^{1}H$ 2D TOCSY spectra with a mixing time of 70 ms showed many signal ridges and distortions and overall a very low signal-to-noise ratio. The 30 ms data is somewhat better, but still only has  limited signals useful for the assignment. Note that for the preparation of bicelles, only deuterated DMPC ($d_{54}$) was used, whereas DMPG, CL and DihepPC were fully protonated. In case of SDS micelles only deuterated SDS ($d_{25}$) was used. Thus, the specta in SDS micelles (Fig. S5) look overall better.

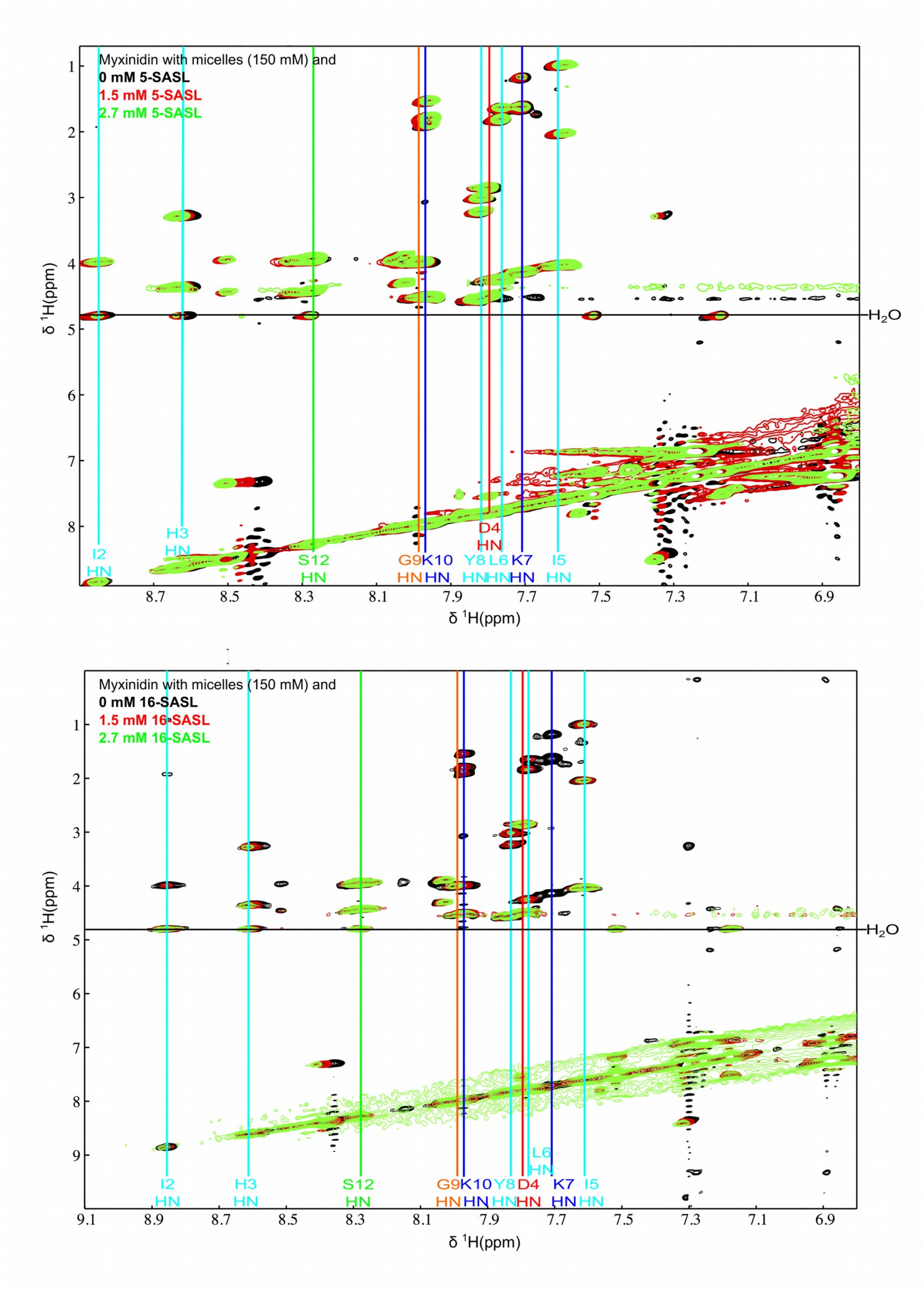


**Fig. S6:** Superposition of the $^{1}H$-$^{1}H$ 2D TOCSY spectra of myxinidin in the presence of negatively charged membrane mimetic SDS micelles in the absence of 5-SASL (top) or 16-SASL (bottom) or increasing amounts of either. The color coding and the spin label concentration are given in the upper left of each plot. The assigned amide and aromatic protons are labeled with the one-letter amino acid code, the residue sequence position and the atom name.

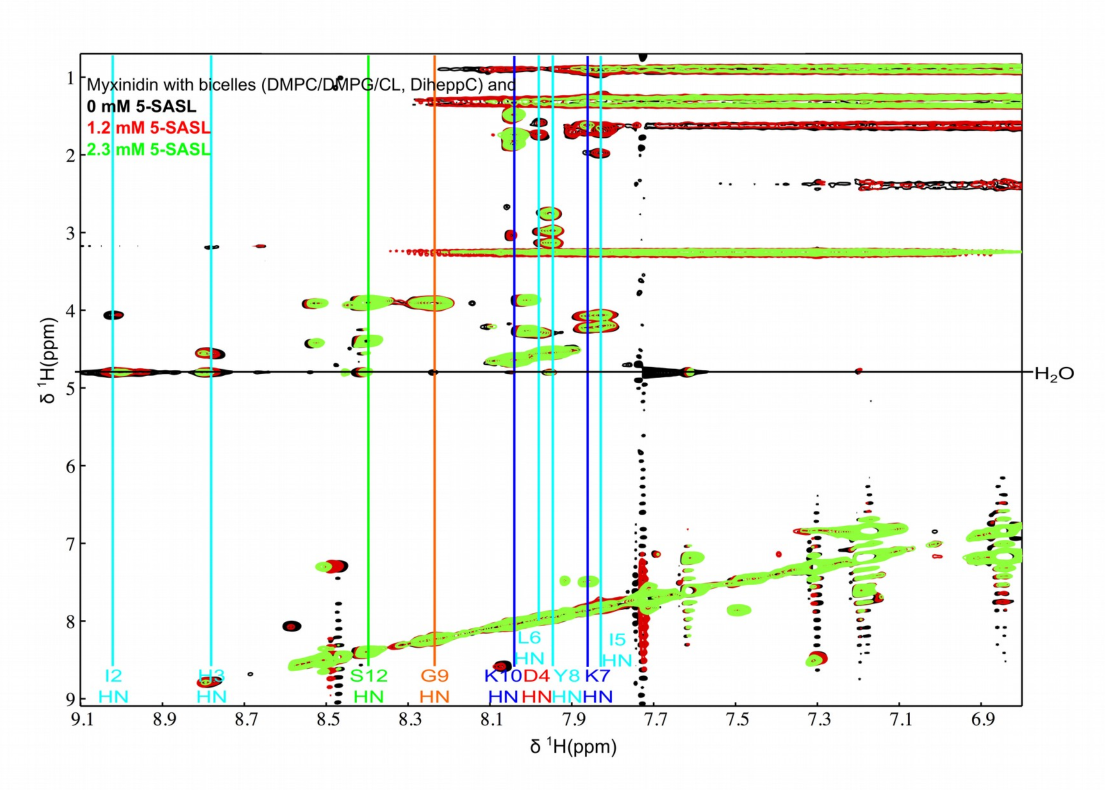


**Fig. S7:** Superposition of the $^1$H-$^1$H 2D TOCSY spectra of myxinidin in the presence of negatively charged membrane mimetic DMPC/DMPG/CL bicelles, both in the absence of 5-SASL and in the presence of increasing amounts of 5-SASL. The color coding and the spin label concentration are given in the upper left corner of each plot. The assigned amide and aromatic protons are labeled with the one-letter amino acid code, the residue sequence position and the atom name.

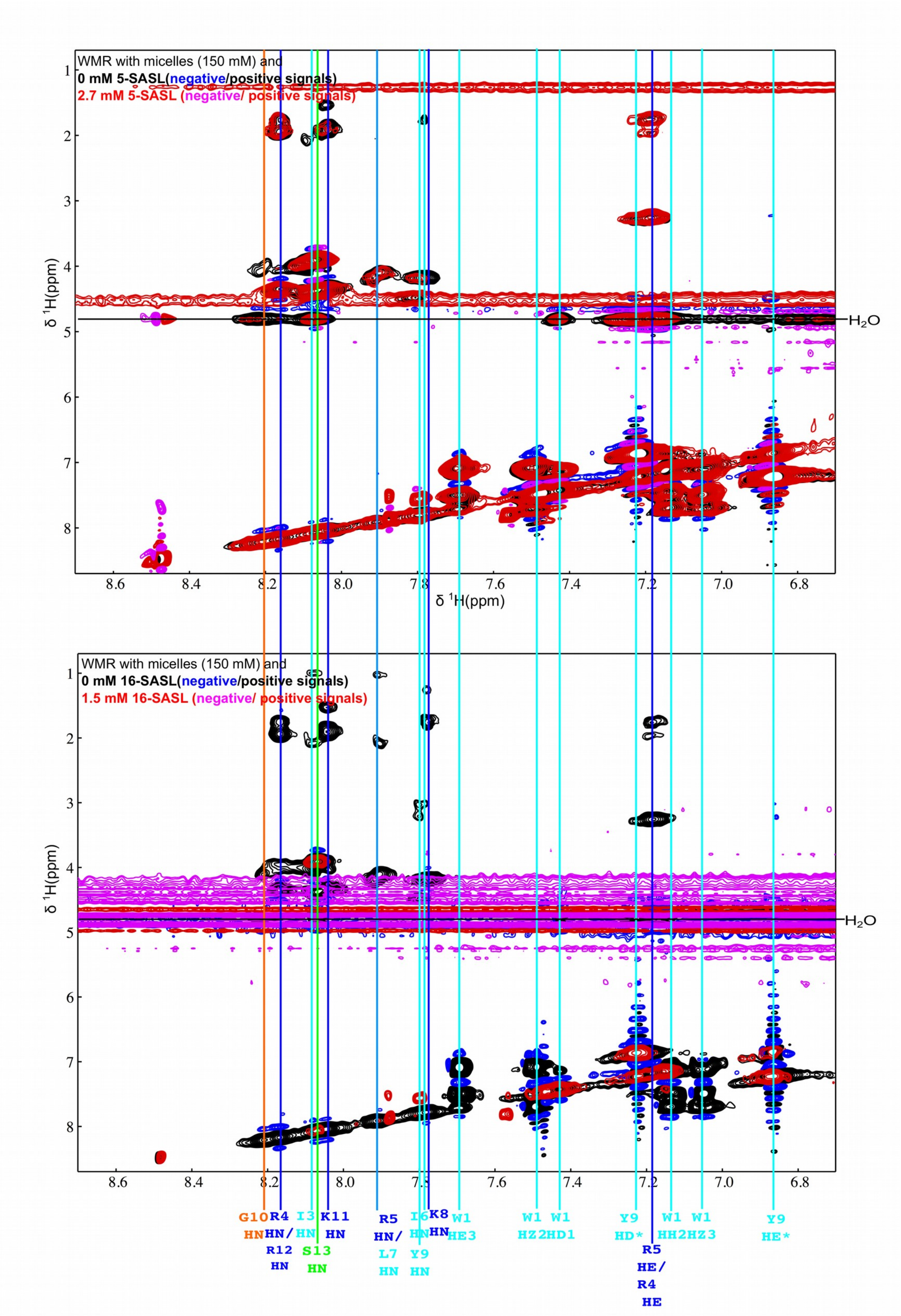


**Fig. S8:** Superposition of the ¹H-¹H 2D TOCSY spectra of WMR in the presence of negatively charged membrane mimetic SDS micelles in the absence of 5-SASL (top)

or 16-SASL (bottom) or increasing amounts of either. The color coding and the spin label concentration are given in the upper left corner of each plot. Due to the signal overlap in the region showing the HN-Hα crosspeaks (about 7.5-8.2 ppm & 3.5-4.5 ppm) as well as signal distortions and a quite strong signal broadening and intensity reduction due the presence of the paramagnetic spin label, the data could not be analyzed quantitatively. The assigned amide and aromatic protons are labeled with the one-letter amino acid code, the residue sequence position and the atom name.

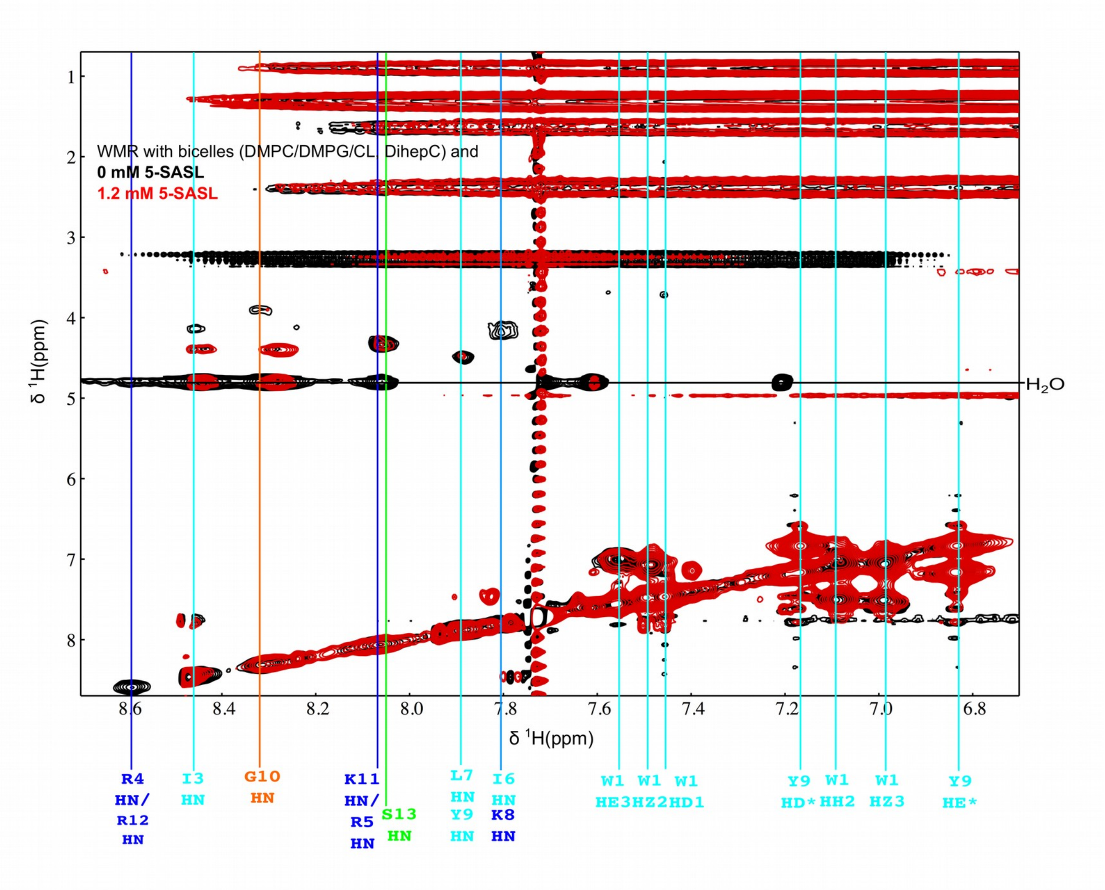


**Fig. S9:** Superposition of the $^{1}H$-$^{1}H$ 2D TOCSY spectra of WMR in the presence of negatively charged membrane mimetic DMPC/DMPG/CL bicelles, both in the absence of 5-SASL and in the presence of increasing amounts of 5-SASL. The color coding and the spin label concentration are given in the upper left corner of each plot. Due to the signal overlap in the region showing the HN-Hα crosspeaks (about 7.5-8.5 ppm & 3.5-4.5 ppm) as well as signal distortions and a quite strong signal broadening and intensity reduction due the presence of the paramagnetic spin label, the data could not be analyzed quantitatively. The assigned amide and aromatic protons are labeled with the one-letter amino acid code, the residue sequence position and the atom name.

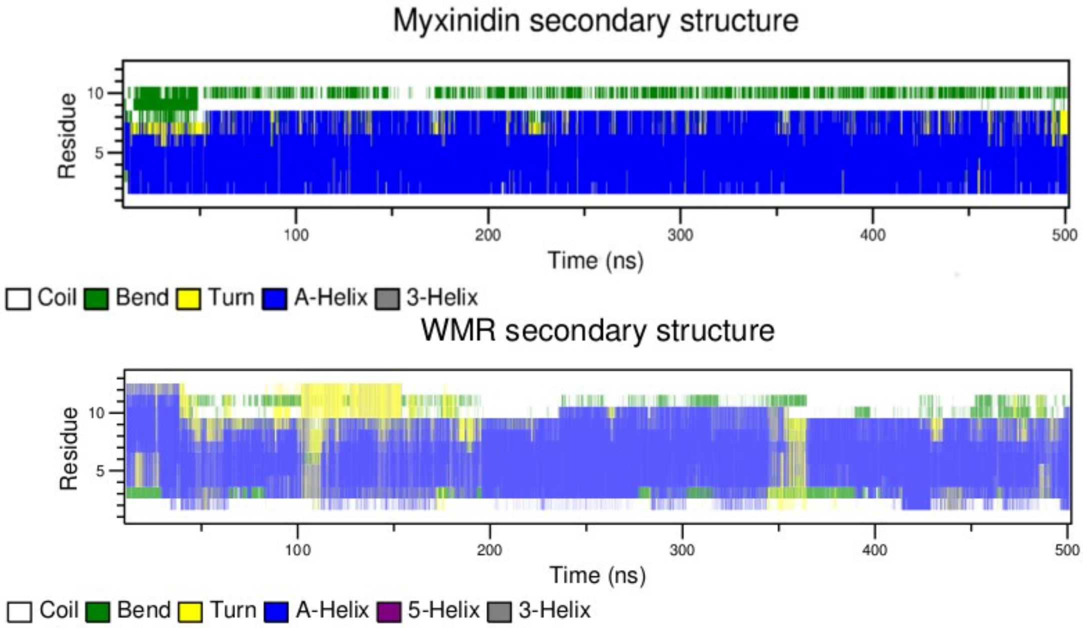


**Fig. S10:** Analysis of the secondary structure content of myxinidin and WMR in the presence of SDS micelle as a function of the simulation time.
